# Model-based characterization of the equilibrium dynamics of transcription initiation and promoter-proximal pausing in human cells

**DOI:** 10.1101/2022.10.19.512929

**Authors:** Yixin Zhao, Lingjie Liu, Adam Siepel

## Abstract

In metazoans, both transcription initiation and the escape of RNA polymerase (RNAP) from promoter-proximal pausing are key rate-limiting steps in gene expression. These processes play out at physically proximal sites on the DNA template and appear to influence one another through steric interactions, leading to a complex dynamic equilibrium in RNAP occupancy of the ~100 bp immediately downstream of the transcription start site. In this article, we examine the dynamics of these processes using a combination of statistical modeling, simulation, and analysis of real nascent RNA sequencing data. We develop a simple probabilistic model that jointly describes the kinetics of transcription initiation, pause-escape, and elongation, and the generation of nascent RNA sequencing read counts under steady-state conditions. We then extend this initial model to allow for variability across cells in promoter-proximal pause site locations and steric hindrance of transcription initiation from paused RNAPs. In an extensive series of simulations over a broad range of parameters, we show that this model enables accurate estimation of initiation and pause-escape rates even in the presence of collisions between RNAPs and variable elongation rates. Furthermore, we show by simulation and analysis of data for human cell lines that pause-escape is often more strongly rate-limiting than conventional “pausing indices” would suggest, that occupancy of the pause site is elevated at many genes, and that steric hindrance of initiation can lead to a pronounced reduction in apparent initiation rates. Our modeling framework is generally applicable for all types of nascent RNA sequencing data and can be applied to a variety of inference problems.

## Introduction

Across all branches of life, homeostatic control of gene expression arises from a series of dynamic equilibria among competing processes. For example, cellular RNA concentrations reflect an equilibrium between RNA production and decay; RNA polymerase (RNAP) occupany reflects an equilibrium among transcription initiation, elongation, and termination; and protein concentrations reflect an equilibrium between protein synthesis and degradation. When such concentrations change, say, across conditions, cell types, or developmental stages, it is typically because the relative rates of competing processes are altered and a new equilibrium is reached, rather than because any individual process is wholly enabled or disabled.

For decades, research on the regulation of eukaryotic gene expression focused on RNAP recruitment and transcription initiation, which were thought to be the rate-limiting steps under most circumstances [1]. In recent years, however, it has become clear that many downstream steps in transcription can be regulated. One striking observation, first noted at particular loci in *Drosophila* and mammals [2–6], is that RNAPs are frequently held in a “paused” position about 30–60 bp downstream of the transcription start site, before escaping into productive elongation [7–14]. Later studies showed that such promoter-proximal pausing is widespread across metazoans [14]. Several lines of evidence indicate that the escape from promoter-proximal pausing is frequently a regulated step in gene expression [15]. Regulation at the pause-escape stage appears to be particularly advantageous when a rapid transcriptional response is required, as in heat shock or other stimulus-controlled pathways [11, 15–17].

It has been noted that promoter-proximal pausing not only has a direct effect on the rate at which RNAPs proceed into productive elongation, but may also have an indirect influence on the rate of productive initiation, owing to steric inference between paused and initiating RNAPs. The key observation is that the physical space immediately downstream of the transcription start site (TSS) is limited: each RNAP has a “footprint” of ~33–35 bp [18,19] and the pause site is typically about ~50 bp downstream of the TSS. Thus, depending on precisely how much space is required between two adjacent RNAPs, there is typically room for only one, or perhaps two or three, RNAPs in the pause region before new initiation events begin to be blocked. Indeed, computer simulations of the movement of RNAPs along the DNA template have suggested that such steric hindrance could substantially reduce initiation rates [19]. More recently, genome-wide studies of human [20, 21] and *Drosophila* [22] cells found strong evidence that initiation rates were restricted by pause-escape rates at many genes. Thus, it appears that rates of productive initiation are often governed by a dynamic equilbrium between transcription initiation and pause-escape.

In recent years, two types of approaches have dominated in the study of transcriptional dynamics: (1) high-resolution imaging approaches based on fluorescence *in situ* hybridization, photobleaching, or electron micrography in single cells (e.g., [23–27]); and (2) genomic approaches based on nascent RNA sequencing (NRS) or chromatin immunoprecipitation (ChIP) and sequencing across populations of cells (e.g., [15, 20, 22]). The imaging approaches allow for more direct characterization of the dynamics of individual RNAPs, but as yet, they cannot be carried out at scale; instead, they are typically applied to one or a few loci. The genomic approaches are more indirect, requiring tagging and pull-down of sequences such as newly synthesized RNA or RNAP-associated DNA. In addition, because they produce summaries for large populations of cells, they typically require fairly complex statistical analyses for interpretation. Nevertheless, these methods have the crucial advantage of being applicable at genome-wide scale for modest cost, by exploiting the many recent advances in genome-sequencing and related technologies.

For these reasons, we focus in this article on the study of transcriptional dynamics using genomic data. We focus in particular on the use of data from NRS methods—such as GRO-seq [9], PRO-seq [28,29], NET-seq [30, 31], and TT-seq [32]—which have matured dramatically in recent years, with major improvements in resolution, background signal, ease-of-use, and cost. These methods can be thought of as producing “snapshots” of the positions of engaged RNAPs across a population of cells either at steady-state or in a time course after a stimulus is applied. We develop a general statistical modeling framework that allows interpretation of these snapshots in a manner that reveals rates of transcriptional initiation, promoter-proximal pause escape, and elongation, as well as interrelationships among these processes. Notably, we focus on modeling transcriptional dynamics under equilibrium conditions, which, in comparison to studies of time courses following transcriptional stimuli, allows us to examine larger sets of genes and avoid the potential off-target effects of commonly used drugs such as triptolide. The focus on steady-state conditions also leads to relatively simple and interpretable mathematical results. Importantly, our methods are applicable not only to newly produced NRS data sets, but to the thousands of sequenced samples that are already publicly available in databases such as the Gene Expression Omnibus [33]. We apply these new methods to both simulated and real NRS data, and refine our model to account for variable pause sites across cells and steric hindrance of transcription initiation from paused RNAP. Altogether, we find strong evidence that promoter-proximal pausing has major importance in the dynamics of transcription in human cells, that RNAP occupancy in the pause region tends to be high at many genes, and that initiation rates are frequently limited by paused RNAPs. We discuss various implications of these findings in detail.

## Results

### A simple probabilistic model for transcription initiation, promoter-proximal pausing, and elongation

Our initial model consists of two layers: a continuous-time Markov model for the movement of individual RNA polymerases (RNAPs) along a transcription unit (TU), and a conditionally independent generating process for the read counts at each nucleotide site (**Fig. 1A&B**) [34]. Together, these components produce a full generative model for NRS read counts along the TU, permitting inference of transcriptional rate parameters from the raw data. The Markov model consists of a state *Z_i_* for each nucleotide position *i* ∈ {1*,…, N*} of the RNAP plus an additional state, *Z*_0_, that represents free RNAPs. It is parameterized by a transcription-initiation rate *α*, a rate of promoter-proximal pause escape *β*, and a termination rate *γ*. In addition, it includes an elongation rate *ζ_i_* for each position *i*, which can either be assumed constant across sites (with *ζ_i_* = *ζ* as throughout this manuscript) or allowed to vary. For mathematical convenience (see **Methods**), the parameters *α*, *β*, and *γ* are multiplied by the corresponding *ζ_i_* parameters. In this manuscript, we focus on inference of *α* and *β* and largely ignore *γ*, because the 3′ ends of TUs are difficult to characterize.

**Figure 1:**
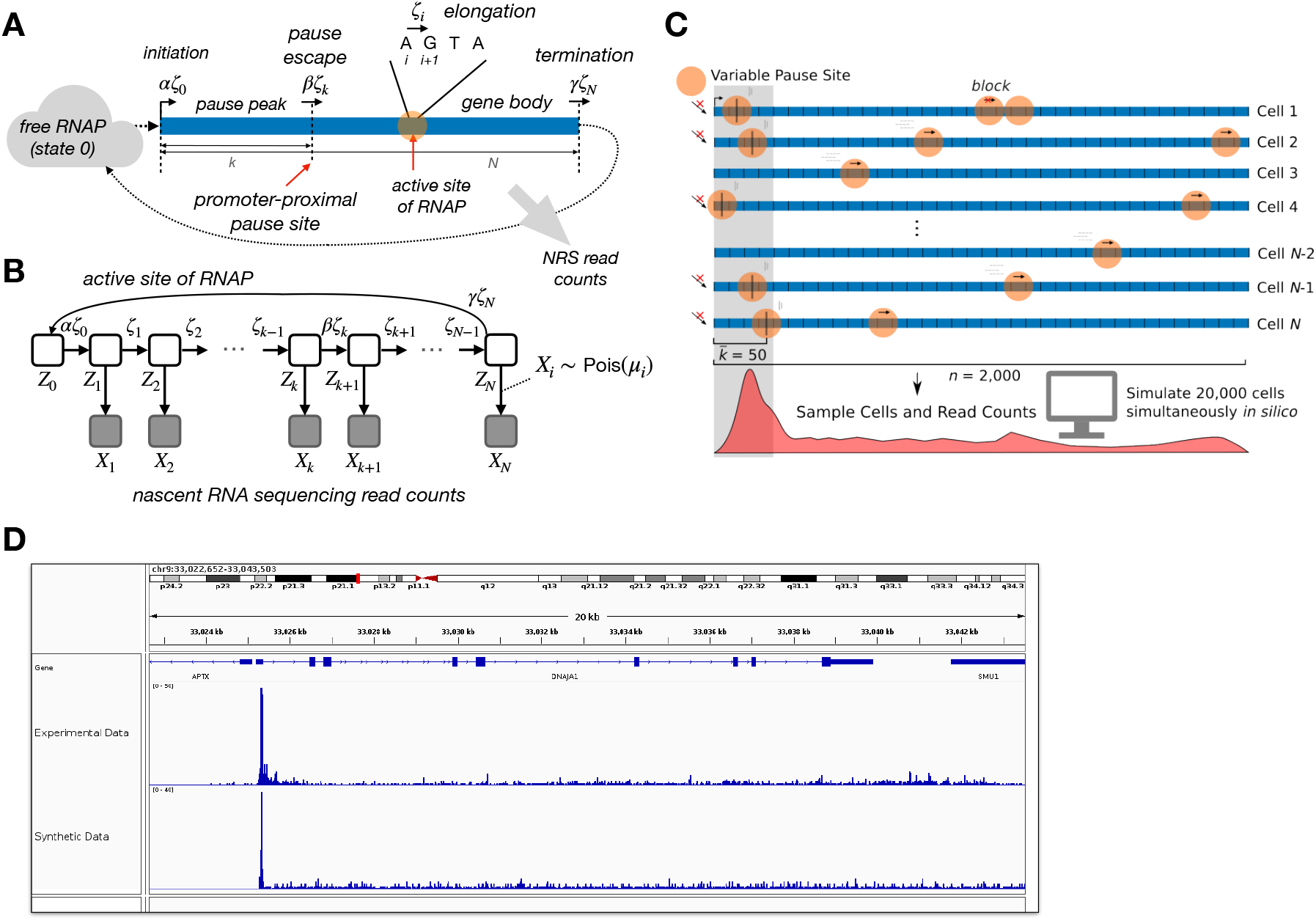
**A**. Conceptual illustration of model, focusing on the kinetic model for RNAP movement on the DNA template. Gray arrow indicates that a second layer of the model describes generation of nascent RNA sequencing (NRS) read counts based on the distribution of RNAP positions across cells. **B**. Graphical model representation with unobserved continuous-time Markov chain (*Z_i_*) and observed read counts (*X_i_*). Read counts at each site *X_i_* are conditionally independent and Poisson-distributed given mean *μ_i_*, which reflects both the density *P* (*Z_i_*) and the sequencing depth *λ*. **C**. Design of SimPol (“Simulator of Polymerases”). Based on user-defined initiation, pause-escape, and elongation rates, SimPol tracks the movement *in silico* of RNAPs across *n*-bp DNA templates in *N* cells, then samples synthetic read counts based on RNAP positions. SimPol identifies collisions and prohibits RNAPs from passing one another. It also models variable pause sites and elongation rates. **D**. Example of synthetic nascent RNA sequencing data from SimPol, shown in IGV [70] alongside matched real PRO-seq data from ref. [36] for the *DNAJ1* gene on chromosome 9 of the human genome.

In this version of the model, promoter-proximal pausing is assumed to occur at a fixed position *k* along the DNA template, where *k* can be pre-estimated from the data. This assumption will be relaxed in subsequent sections. We further assume that the read counts at each position *i* are Poisson-distributed with mean *μ_i_*, where *μ_i_* is a scaled version of the probability density of *Z_i_* (and, hence, the RNAP density at nucleotide *i*) that reflects the initiation and elongation rates, as well as the sequencing depth. Finally, in order to make use of a time-homogeneous Markov chain, we assume that RNAPs are sufficiently sparse along the DNA template that collisions between them are rare—another assumption that will be relaxed later. Notably, the model can be applied either in a nonequilibrium setting based on time-course data or to a single data set at steady state. We focus in this manuscript on the steady-state case, which is more mathematically convenient and interpretable, yet, as we show below, still reveals a great deal about the dynamics of transcription. In order to ensure identifiability of initiation and pause-escape rates at steady state, we assume that initiating RNAPs remain on the DNA template for its entire length, with negligible rates of premature termination (see **Discussion**).

This simple version of the model results in convenient, closed-form maximum-likelihood estimators for *β* and a parameter closely related to *α*. Because the initiation rate *α* is confounded at steady-state with the elongation rate *ζ* and the sequencing depth *λ*, we instead work with a compound parameter 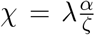 representing the read-depth-scaled ratio of the initiation rate to the elongation rate. It turns out that the MLE for *χ* is simply equal to the average read depth along the gene body and the MLE for *β* is given by the ratio of the average read depth in the gene body to that in the pause peak (see **Methods**). The estimates of both *α* and *β* can be considered *relative* to *ζ*, based on the multiplicative parameterization of the model. Notably, similar average-read-depth estimators have been widely used in the analysis of nascent RNA-sequencing data, typically with more heuristic justifications. For example, the inverse of the estimator for *β* is commonly known as the “pausing index” [14, 35], and the estimator for *α* is often used as a general measure of transcription output [17]. In our case, these estimators emerge as MLEs under a generative probabilistic model, making it possible to characterize the dynamics of transcription from raw NRS read counts at steady state.

To examine the quality of these estimators, we developed a flexible computational simulator, called SimPol (“Simulator of Polymerases”), that tracks the progress over time of individual RNAPs across DNA templates in thousands of cells under user-defined initiation, pause-escape, and elongation rates (**Fig. 1C**). Unlike our assumed model, the simulator tracks potential collisions between RNAPs and prohibits one RNAP from passing another along the template. In addition, the simulator allows for variation across cells in pause-site location and variation across both cells and nucleotide positions in local elongation rate (see **Methods** for details). After running to equilibrium, the simulator generates synthetic read counts at each nucleotide position by sampling the positions of RNAPs across cells, and then sampling read counts conditional on local RNAP frequency.

Using SimPol, we generated synthetic data sets for a range of plausible initiation and pause-escape rates, and read depths corresponding median expression levels from real data (see **Methods**). We performed separate sets of simulations for cases where the pause site is fixed at a known value *k* and for cases where *k* is allowed to vary across cells according to a truncated Gaussian distribution. We assumed a mean value of *k* equal to 50 bp and a standard deviation of 25 bp, approximately as we observe in real data (see **Methods**), and required a minimum center-to-center distance between adjacent RNAPs of 50 bp (as in [20]). For each simulated data set, we estimated *χ* and *β* from the synthetic data and compared the estimated and true rates. In cases where the pause site was variable, we estimated *β* by using the average read depth across a 200 bp pause peak region, similar to other studies that have used a pausing index (e.g., [17]). In addition, for *β*, we evaluated the product *βζ*—which represents the absolute rate of pause-escape—assuming the mean value of *ζ* = 2 kb/min that was used for simulation. For *α*, we calibrated our estimates relative to a baseline case simulated with *αζ* = 1 and no pausing and again assumed *β* = 2 kb/min (see **Methods**). In this way, we were able to account for both the sequencing read depth and the elongation rate, and express both the initiation and pause-escape rates in absolute units of events per minute, for ease of interpretation.

We found that these initial estimates of *α* and *β* were accurate in some cases but significantly biased in others (**Fig. 2**, **Supplementary Fig. S1**). In particular, estimates of the initiation rate *α* were close to the truth when *α* was not too large and *β* was not too small; but either a moderately high *α* or a moderately low *β* led to a notable downward bias in the *α* estimates (**Fig. 2A**). This bias ranged from ~50% when *α* was high or *β* was low but the other parameter was in the favorable regime, to more than an order of magnitude when both parameters were unfavorable. As we explore below, these biases suggest an influence from steric hindrance of new initiation events owing to RNAPs in the pause peak, which would be expected to lead to under-estimation of *α* precisely when pause-escape rates are low and/or initiation rates are high. Importantly, these biases can be pronounced for plausible values of *α* and *β*, suggesting that it can be misleading to treat average read counts in the gene body as a measure of the initiation rate (see **Discussion**).

**Figure 2:**
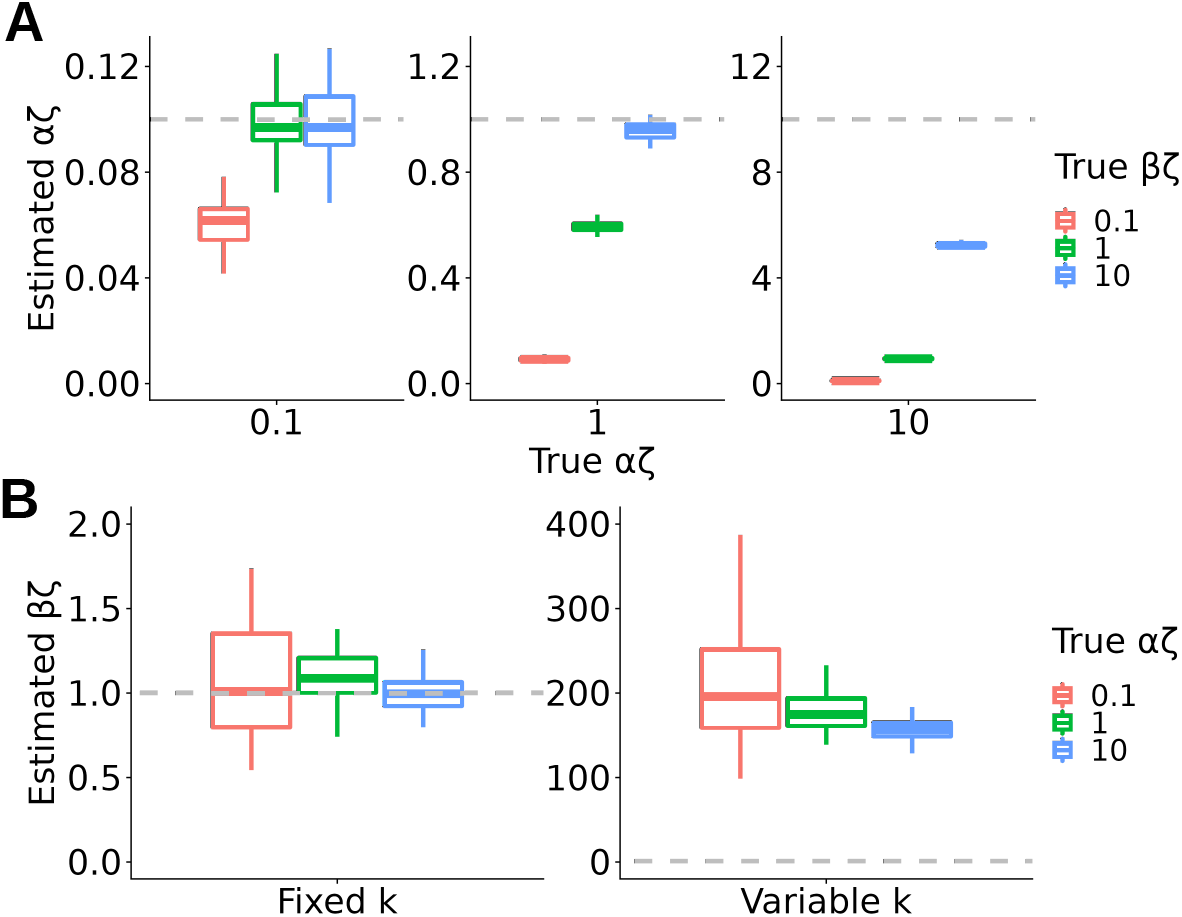
Accuracy of estimated values of the transcription initiation rate *α* and pause-escape rate *β* under the initial version of the model. Estimates are expressed as products with the elongation rate *ζ* (*αζ* and *βζ*). **A**. Simulated true vs. estimated values of *αζ*, for *αζ* ∈ {0.1, 1, 10} (left to right) and *βζ* ∈ {0.1, 1, 10} (see key). Dashed lines indicate the ground truth. **B**. Estimated values of *βζ* for simulated true values of *βζ* = 1 and *αζ* ∈ {0.1, 1, 10} (see key), when the pause-site *k* is fixed (left) or variable across cells (right) in simulation. Dashed lines indicate the ground truth. Results for other values of *βζ* are shown in **Supplementary Fig. S1**. All boxplots summarize 50 replicates of the simulation; box boundaries indicate 1st and 3rd quartiles, and horizontal line indicates median. A value of *ζ* = 2 kb/min is assumed; *αζ* and *βζ* can be assumed to have units of events per minute. Pause sites occur at a mean position of *k* = 50 bp. In the variable case, we assume a Gaussian distribution with a standard deviation of 25 bp.

By contrast, in the case where the pause site was held fixed and assumed known during estimation, estimates of *β* were accurate across a range of true *α* and *β* values (**Fig. 2B**, left side; **Supplementary Fig. S1**), indicating that the model describes the dynamics of pause-escape well. When the distance to the pause site, *k*, was allowed to vary across cells, however, and the read-depth in the pause peak was estimated by averaging across sites, the increased read density owing to pausing was dramatically under-estimated, resulting in a strong upward bias in estimates of *β* (**Fig. 2B**, right side; **Supplementary Fig. S1**). In this case, the “spike” of read counts becomes a rounded “peak” of reduced height, which naturally biases the estimate of *β*. As it turns out, this simple source of bias is deceptively difficult to eliminate. For example, it is possible to mitigate it by using the maximum rather than the average read depth in the pause peak, but it remains fairly pronounced. Instead, we develop an extension to the model to remedy this problem in the next section.

### Promoter-proximal pause sites vary across cells and pause-escape is rate-limiting

To address the bias in estimation of *β* in the presence of variable pause sites, we extended the model to allow for a distribution of pause sites across cells (see **Methods**). In this version of the model, the read counts at each site in the pause peak are assumed to arise from a mixture of cells in which an RNAP is and is not paused at that position. If a Gaussian distribution of pause sites is assumed, the model can be fitted to the data relatively simply and efficiently by expectation maximization (EM). This procedure results in maximum-likelihood estimates not only of *χ* and *β* but also of the mean and variance of the pause-site position across cells. Thus, this version of the model no longer requires prior information about the pause-site *k*, but instead allows its mean and variance to be estimated separately at each gene from the raw data.

We reanalyzed our simulated data using this version of the model and found that it was highly effective at correcting the bias in estimated values of *β*. In this case, the same model addresses the fixed and variable pause-site scenarios equally well across a range of values of *α* and *β* (**Fig. 3A**; **Supplementary Fig. S2**). Whereas the averaging approach described in the previous section resulted in over-estimates of *β* by more than two orders of magnitude, that bias was completely eliminated when variability in the pause site was modeled directly. Thus, typical average-based estimates of a “pausing index” (*I_P_*) for real data may substantially under-estimate the prominence of promoter-proximal pausing in RNAP dynamics (see **Discussion**). The model addresses this problem. Only when *α* is quite small and *β* is quite large (bottom of **Supplementary Fig. S2**), resulting in few excess reads in the pause peak, do the model-based estimates sometimes exhibit a substantial downward bias. Notably, while the full model requires an iterative method such as EM, it is possible to derive an approximate closed-form expression for *β* in terms of the pausing index *I_P_*, which clarifies the physical meaning of *I_P_* (see **Methods** and **Discussion**).

**Figure 3:**
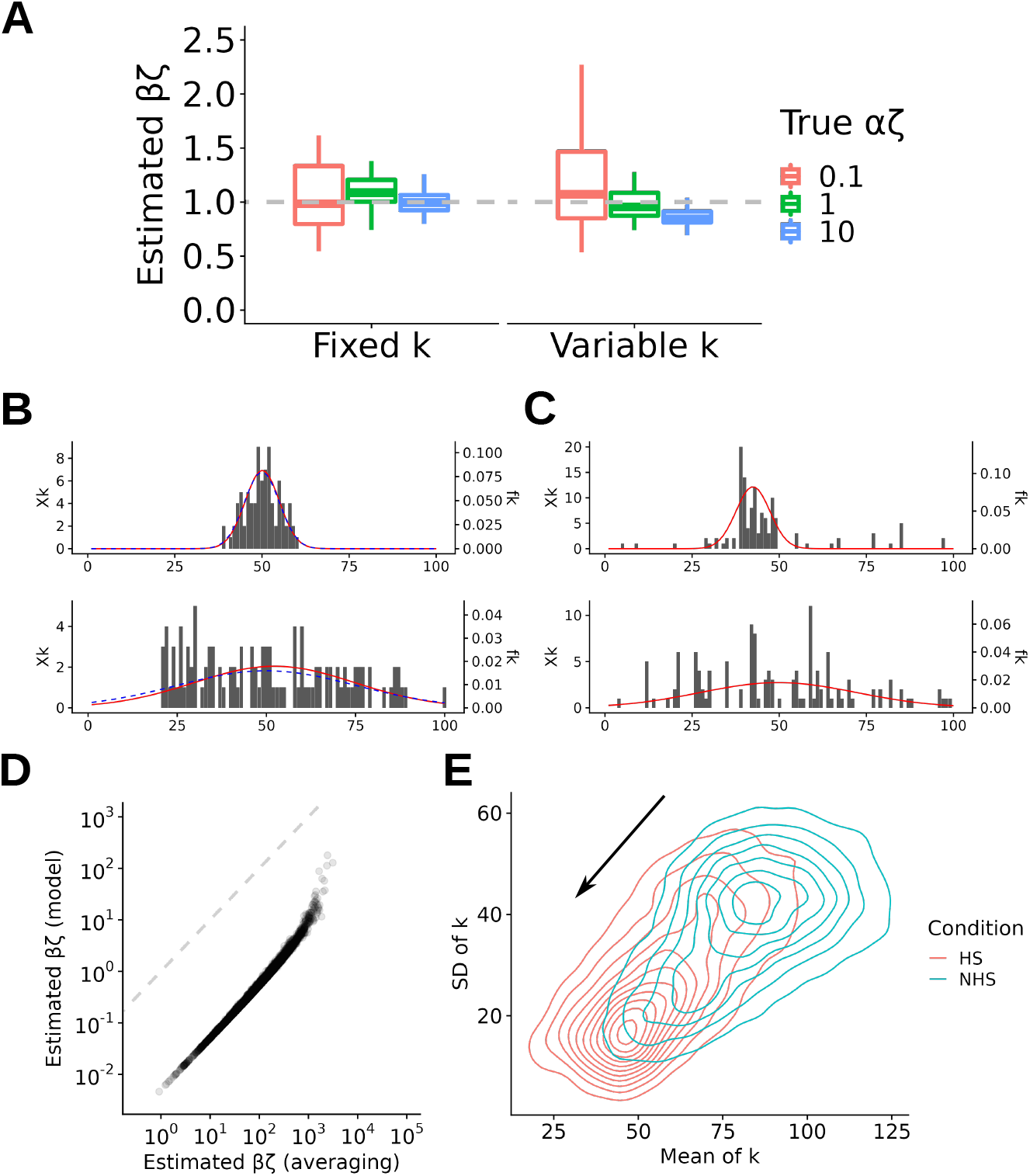
**A**. Accuracy of estimated values of the pause-escape rate *β* under the version of the model that allows for a distribution of pause-sites *k* across cells. Shown are estimated values of *βζ* for simulated true values of *βζ* = 1 and *αζ* ∈ {0.1, 1, 10} (see key), when the pause-site *k* is fixed (left) or variable across cells (right) in simulation. Dashed lines indicate the ground truth. Results for other values of *βζ* are shown in **Supplementary Fig. S2**. All boxplots summarize 50 replicates of the simulation; box boundaries indicate 1st and 3rd quartiles, and horizontal line indicates median. Simulated pause sites occurred at a mean position of *k* = 50 bp. In the variable case, we assumed a Gaussian distribution with a standard deviation of 25 bp. **B**. Examples of pause peaks in simulated data, showing assumed distribution of pause sites (blue dashed line) and distribution inferred by expectation maximization (red solid line). **C**. Similar examples from real data from ref. [36]. **D**. Estimates of *βζ* under the original averaging approach (horizontal axis) vs. estimates of *βζ* under the model that allows for variable *k* across cells (vertical axis). **E**. Contour plot showing the distribution of estimated means (horizontal axis) and standard deviations (vertical axis) of the pause peak position *k*, under the “no heat shock” (NHS) and “heat shock” (HS) conditions. Data from ref. [36]. In panels A and D, a value of *ζ* = 2 kb/min is assumed; thus, *αζ* and *βζ* can be assumed to have units of events per minute.

To validate the ability of our model to capture differences in the pause-site distributions across cells, we applied it to synthetic data sets with larger and smaller variances in pause-site locations. We found that the model was able to recover the correct distributions fairly accurately, as long as the read counts in the pause peak were not too sparse (**Fig. 3C**). Notably, the model is provided with no prior information about the mean or variance in the pause-site location, but is only constrained to consider a range of possible values (here from *k*_min_ = 1 to *k*_max_ = 200). When applied to real data, the model also appears to do a good job of identifying plausible distributions across cells in the locations of pause sites (**Fig. 3D**).

Having observed good performance on synthetic data, we applied the model to published PRO-seq data for K562 cells before and after heat shock [36], focusing at first on the untreated sample. For comparison, we also fitted the previous version of the model (based on the average read depth in the pause peak) to the same data set. We found, as expected, that the distribution of *β* estimates was substantially shifted, with values about two orders of magnitude smaller under the new model (**Fig. 3D**). Based on our simulation results, we expect these smaller estimates to much more accurately represent the true pause-escape rates.

For comparison with experimental results (e.g., [20, 37]), we converted these estimated pause-escape rates to half-lives for RNAP residence in the pause region, conditional on an assumed elongation rate of *ζ* = 2 kb/min [16, 20, 37]. For a direct comparison, we considered the time required for elongation up to the pause site, as well as the waiting time for escape (see **Methods**). We estimated a median half-life of 1.4 min. and a mean of 2.8 min. (**Supplementary Fig. S3A**). These estimates are fairly similar to ones obtained experimentally by Gressel et al. [20] for CDK9-inhibited human Raji B cells (median pause duration of 1 min, corresponding to a median half-life of 0.7 min), but somewhat smaller than those from some other recent experimental studies (e.g., [37]; see **Discussion**). We additionally examined PRO-seq data from another study of K562 cells [17] and estimated a similar distribution of half-lives, with a median of 1.2 min. and mean of 2.1 min. (**Supplementary Fig. S3B**). Notably, induction of heat shock and treatment with the drug celastrol both increased half-lives by 2–3-fold (**Supplementary Fig. S3C&D**), consistent with the observations of increased pause indices in the original reports [17, 36].

Interestingly, the pause peaks estimated for both of these data sets varied substantially across genes in their breadth, as quantified by the estimated variance across cells in the pause-site position 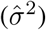. Hypothesizing that the underlying DNA sequences might contribute to the precision of pausing, we searched for sequence motifs that distinguished the pause regions having the narrowest peaks (10% with smallest 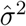) from those having the broadest peaks (10% with largest 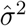), focusing again on the untreated samples (see **Methods**). In both the heat-shock [36] and celastrol [17] data sets, we found that the most strongly enriched motif in the narrow peaks closely matched the binding motif for TATA-Box Binding Protein Associated Factor 1 (TAF1), the largest subunit of general transcription factor IID (TFIID) and a key component of the pre-initiation complex (PIC) [38, 39] (**Supplementary Figs. S4 & S5**, panels **A & B**). Consistent with these motif enrichments, chromatin immunoprecipitation and sequencing (ChIP-seq) data for TAF1 in K562 cells from the ENCODE project [40] exhibits a substantially stronger signal at narrow peaks than at broad peaks, particularly near the TSS (**Supplementary Figs. S4C & S5C**). Interestingly, it has recently been shown that the PIC alone is sufficient to establish RNAP pausing, and that rapid TAF1 depletion induces pause-release genome-wide [41]. We found other enriched motifs, but they mostly consisted of repetitive, highly G+C-rich sequences, similar to observations in other recent studies [20, 42, 43]. We examined the DNA sequences surrounding the nucleotide with the maximum PRO-seq read count in each pause peak, and observed a clear enrichment for cytosines at this position (see also [20, 42–44]; **Supplementary Figs. S4D & S5D**). However, this enrichment was evident in both the narrow and broad peaks.

We were also interested in possible effects of cellular stress on pause-site locations. Metaplots summarizing the accumulated PRO-seq signal across large classes of genes have suggested that pause peaks may tend to shift in location and become sharper following stress, particularly for downregulated genes (e.g., [29, 36]). To test whether our model supported such a change, we compared our estimates of the pause site locations, before and after application of heat shock (**Fig. 3E**). Indeed, we observed a striking reduction in both the mean and variance of *k* after heat shock. This shift may reflect differences in activity of protein complexes required for pausing, including NELF, DSIF, P-TEFb, or the PIC (e.g., [41,45]). For comparison, we examined PRO-seq data from the celastrol study, which reported a stress response that resembles heat shock in some ways [17]. In this case, however, we did not observe a clear change in the mean and variance of *k* (**Supplementary Fig. S6**).

### High occupancy at promoter-proximal pause sites hinders transcription initiation

To test the hypothesis that the observed under-estimation of *α* is driven by steric hindrance of initiation (**Fig. 2A**), we collected two types of auxiliary data from our simulation experiments. First, we tracked the fraction of cells in which the “landing pad” for a potential new initiation event was already occupied by an RNAP (“landing-pad occupancy”), which is a close proxy in our simulations for the fraction of potential initiation events that were not allowed to occur owing to steric hindrance. We found that this fraction was frequently quite high (often 50% or more, and in some cases ≥90%), and tended to be highest when the bias in estimated *α* was most pronounced (**Supplementary Fig. S7A**). Second, we measured the rate at which initiation events successfully occurred (not being blocked) and compared it with our estimates of *α*, finding much better agreement between this “effective” (sometimes called “productive” [20]) initiation rate and our estimates (**Supplementary Fig. S7B**). Finally, we found that our estimation accuracy for *α* (as measured by the ratio of the estimated to true values) had a close negative correlation with the landing-pad occupancy (**Supplementary Fig. S7C**). Together, these findings indicate that—at least in simulation—new initiation events are frequently blocked by RNAPs occupying the region just downstream of the TSS, and that this phenomenon explains much, if not all, of the bias in estimation of *α*.

To address this problem, we extended our probabilistic model to explicitly allow for steric hindrance of initiation at steady state. Briefly, we introduced a distinction between an effective initiation rate *ω* and a potential initiation rate *α*, letting *ω* = (1 − *ϕ*)*α*, where *ϕ* is the landing-pad occupancy. The model assumes that a fraction *ϕ* of new initiation events are blocked by RNAPs occupying the landing pad, and the remaining fraction 1 − *ϕ* are allowed to proceed (see **Methods** for complete details). It allows for more than one RNAP per pause region, because the landing pad will likely be blocked in some cases by RNAPs stacking up behind the pause site. Based on our observations above, we reinterpreted our previous estimator for *α* instead as an estimator for the effective rate *ω*. This new model allows us to compute *ϕ* in terms of *α*, *β*, *k*, and an assumed minimum spacing between RNAPs, denoted *s*. An extended EM algorithm allows joint estimation of *ϕ*, *α*, and *β* from the data, with a Beta prior distribution to ensure that *ϕ* remains in the allowable range, *ϕ* ∈ (0, 1) (**Methods**). In effect, this strategy allows us to correct the assumed model for steric hindrance, yielding estimates not only of the effective initiation rate *ω* and pause-escape rate *β*, as in the previous case, but also of the landing-pad occupancy *ϕ* and, hence, the potential initiation rate *α*.

When we applied this new model to our simulated data, we found that it generally appeared to work as intended. The estimates of *ϕ* were broadly consistent with an empirical measure of the landing-pad occupancy across a range of values of *α* and *β* (**Fig. 4A**), Not surprisingly, given the indirect information about *ϕ* in the data, the variance in these estimates was substantial; nevertheless, their mean values were close to the truth when *ϕ* was small and remained within ~20% of the truth even for large *ϕ*, although the Beta prior did produce a downward bias when *α* » *β* and *ϕ* approached the upper bound of one. The estimates of the pause-escape rate *β* remained excellent overall even in the presence of substantial amounts of steric hindrance (**Supplementary Fig. S8**).

**Figure 4:**
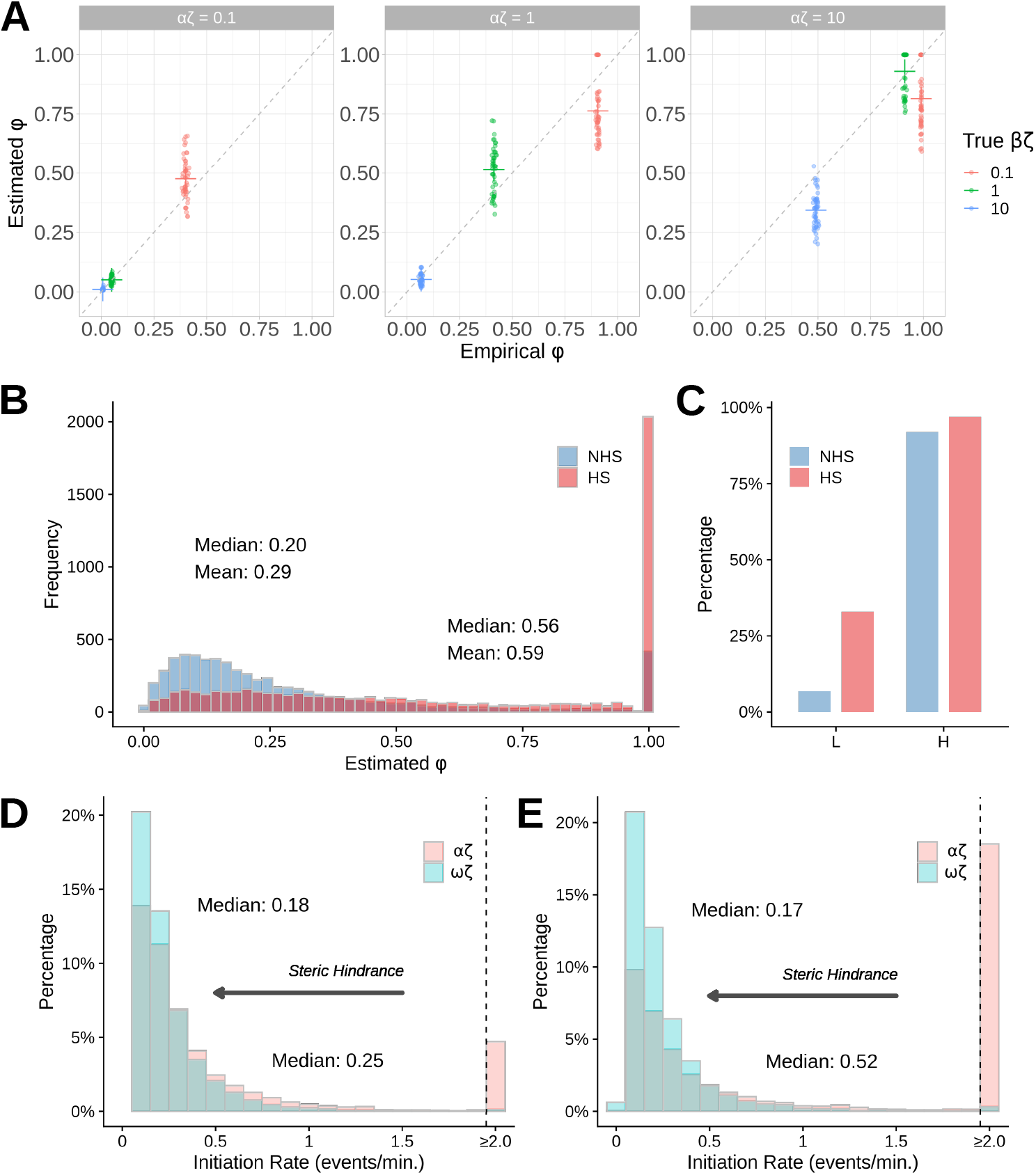
**A**. Accuracy of estimated landing-pad occupancy *ϕ* under the version of the model that allows for steric hindrance in initiation and multiple RNAPs per pause region. Scatter plots show the fraction of simulated cells for which the first 50 bp (the “landing-pad”) are occupied by an RNAP at steady state (“Empirical *ϕ*”) vs. the fraction predicted to be occupied under the model (“Estimated *ϕ*”) based on the simulated NRS data, assuming a minimum spacing of *s* = 50 bp. Results are shown for simulated true values of *αζ* ∈ {0.1, 1, 10} (left to right) and *βζ* ∈ {0.1, 1, 10} (see key), with 50 simulations per parameter combination. Dashed line indicates *y* = *x*, and colored crosses represent the means of the corresponding points. A value of *ζ* = 2 kb/min is assumed, so that *αζ* and *βζ* are in events per minute. **B**. Distribution of estimated *ϕ* for 6,087 robustly expressed genes in K562 cells before (NHS) and after (HS) heat shock under the low (L) calibration [36] (see **Methods** for details). **C**. Percentages of genes having fully occupied landing-pads (*ϕ* ≈ 1) before (NHS) and after (HS) heat shock, under the low (L) and high (H) calibrations. **D & E**. Distributions of scaled estimates of the “effective” (*ωζ*) and “potential” (*αζ*) rates of transcription initiation, in events per minute per cell, for the same genes. Panel **D** represents the NHS case and panel **E** represents the HS case. The *x*-axes are truncated to highlight the bulk of the distributions. Gray arrows indicate effects of steric hindrance.

The most difficult parameter to recover was the potential initiation rate *α*, about which the data are only weakly informative via the effective initiation rate *ω* and the landing-pad occupancy *ϕ*. Nevertheless, *α* was estimated reasonably well when *α* ≤ *β* and *ϕ* ≤ 0.5 (**Supplementary Fig. S9**). When *α* » *β* and *ϕ* → 1, however, the denominator of the estimator 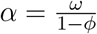 becomes unstable and it is no longer possible to estimate *α* accurately. In these cases, the effective initiation rate is still estimated well, but it is not possible to extrapolate from it to the potential rate, because only a small fraction of potential initiation events are allowed to occur.

Our initial experiments assumed a value of *s* = 50 bp for the minimum center-to-center spacing between adjacent RNAPs (following [20]), but the true value of *s* is not known with any certainty; therefore, we examined the robustness of our model to other plausible values of this parameter. In general, smaller values of *s* will increase the chances that a single paused RNAP will not block the landing pad, but will also allow more RNAPs to stack up behind the pause site. In contrast, when *s* grows larger and approaches *k*, the distance to the pause-site, at most one RNAP can occupy the pause region at a time, and that RNAP will necessarily block the landing pad (see **Methods** for details). We used SimPol to generate data sets for two alternative choices of *s*: a more generous value of *s* = 70 bp and the minimum possible value of *s* = 33 bp. The case of *s* = 33 bp is probably unrealistic given the physical space required for the other components of the PIC (e.g., [41, 46]) but is nevertheless useful as a bound. In our simulations, where *k* has a mean of 50 and standard deviation of 25, ≥ 2 paused RNAPs are possible at 82%, 54%, and 22% of cells for *s* = 33, *s* = 50, and *s* = 70 bp, respectively; and ≥ 3 paused RNAPs are possible at 29%, 2.5%, and 0.02%, respectively.

As expected, we found that our estimates of *ϕ* were best in the case of *s* = 70 bp, when multiple RNAPs per pause region are fairly rare and ≥ 3 almost never occur (**Supplementary Fig. S10**). Notably, however, the case of *s* = 70 bp was only slightly better than the case of *s* = 50 bp, suggesting that our model can accommodate multiple RNAPs at moderate frequency without much error. By contrast, we did see a more noticable bias in the case of *s* = 33 bp, particularly a tendency to over-estimate *ϕ* when *α* = *β* (**Supplementary Fig. S10**). Nevertheless, the estimates of *β* and *ω* remained accurate in all cases.

Using this model we re-analyzed the untreated K562 sample from Vihervaara et al. [36], focusing on 6,087 robustly expressed genes (the top 80% by 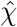). When fitting the model to real data, however, a problem of scale arises: the effective initiation rate *ω* can only be estimated up to a scale factor, which is determined by the (unknown) ratio of the read depth to the elongation rate, *λ/ζ*. At the same time, the pause-release rate *β*—whose estimator is a ratio of two summary statistics (equations 7 and 11)—has a fixed scale; thus, the estimator for the landing-pad occupancy, 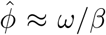(equations 20 and 25), depends on the choice of scale for *ω*.

To address this problem, we calibrated the scale of *ω* using published estimates of the initiation rate. Because there is considerable uncertainty about this quantity in the literature, we selected two different calibration points, one on the low end of the reported range for eukaryotes, at 0.2 events per minute [47–50], and another on the high end, at ~1 event/min (median of estimates reported by [20]). We used a set of “housekeeping” genes [51] for calibration, reasoning that these genes would minimize sensitivity to differences among species, cell types, and conditions. We scaled our estimates of *ω* such that the median value of *ωζ* within housekeeping genes in our set was either equal to 0.2 (the *low* (L) calibration) or equal to the median value in matched housekeeping genes from ref. [20], which was 2.4 events/min (the *high* (H) calibration; see **Methods** for details).

In the case of the L calibration, we found that the distribution of *ϕ* estimates across genes spanned a fairly broad range, with a median of 0.20 and a mean of 0.29 (**Fig. 4B**), suggesting that, on average, almost a third of landing pads are occupied at steady state, but considerable fractions of genes have substantially lower or higher occupancies. Notably, 6.8% of genes were predicted to have their landing-pads fully occupied (*ϕ* ≈ 1; **Fig. 4C**). The scaled estimates of the effective initiation rate *ωζ* were roughly exponentially distributed, with a median of 0.18 and a mean of 0.25 events per minute per cell (**Fig. 4B**). By combining our estimates of *ϕ* and *ωζ*, we were additionally able to obtain per-gene estimates of the potential initiation rate *αζ*. These estimates were modestly inflated, with a median of 0.25 and ~5% of values >2.0 events per minute (**Fig. 4D**). (In most of the cases >2.0, a precise estimate of *αζ* could not be obtained, because *ϕ* approached its limiting value of 1.) Thus, steric hindrance appears to result in a clear reduction in the effective initiation rate in these cells, particularly at a subset of genes having high landing-pad occupancy.

When we assumed the H calibration, the *ϕ* distribution shifted accordingly to the right. Indeed, in this case 92% of genes were predicted to have fully occupied landing pads (*ϕ* ≈ 1; **Fig. 4C**), and as a result, the majority of *αζ* values became undefined, suggesting that this initiation-rate calibration is perhaps too aggressive. Nevertheless, by considering a broad range of potential initiation-rate calibrations, we estimated that the landing-pad occupancy is generally fairly substantial, with the median value of *ϕ* ranging from ~30% to full occupancy. Thus, our model predicts that steric hindrance has a substantial impact on initiation rates at steady state, even in the untreated condition.

For comparison, we repeated the analysis with the treated sample, following heat shock (HS). Consistent with observations of increased pausing after HS [36], we found a dramatic shift toward larger *ϕ* estimates in this sample (**Fig. 4B**). With the L calibration, the fraction of genes with fully occupied landing pads (*ϕ* ≈ 1) increased from 6.8% to 32.9% (**Fig. 4C**). Accordingly, steric hindrance was predicted to have a substantially stronger impact on the effective initiation rate after HS, decreasing from a median of 0.52 events/min to 0.17 events/min (**Fig. 4D**). We also analyzed untreated and treated samples from the K562/celastrol study [17] using 5,964 genes with initiation rates calibrated using similar L and H strategies (**Methods**). The results were qualitatively similar to those from the heat-shock analysis, but the absolute *ϕ* estimates were somewhat lower, shifting from a median of 0.16 in the untreated case to 0.73 after treatment (L calibration; **Supplementary Fig. S12A**). Application of celastrol resulted in a striking increase in fully occupied landing-pads, from 3.3% to 39.4% of genes under the L calibration, but as in the heat shock analysis, the H calibration indicated nearly complete landing-pad occupancy before treatment and a limited shift after treatment (**Supplementary Fig. S12B**). Prediction of *αζ* indicated a modest but clear reduction in initiation rates from steric hindrance before treatment (**Supplementary Fig. S12C**) and a more pronounced one after treatment (**Supplementary Fig. S12D**). Overall, we found that steric hindrance has a clear impact on productive initiation, and that impact is particularly striking during responses to cellular stress.

## Discussion

The widespread occurrence of promoter-proximal pausing has been one of the major surprises of the past ~15 years in the study of gene regulation. Most studies of this phenomenon have focused on its molecular mechanisms and its direct impact on rates of productive elongation [15]. In addition, however, there have been indications that such pausing, when sufficiently pronounced, also imposes indirect limits on rates of transcription initiation [19–21]. In this article, we have shown through analysis of simulated and real data that many aspects of this complex interplay between initiation and pause-escape can be described by relatively simple probabilistic models. Our models not only describe raw read counts as a function of variable rates of initiation and pause-release, enabling estimation of these rates from NRS data, but they also allow for variable pause sites across cells, for reductions in initiation rates via steric hindrance, and for the effects of multiple RNAPs stacking up behind the pause site. We have shown by simulation that both variable pause sites and steric hindrance can have major impacts on the estimated rate parameters. Our analysis of real data indicates not only that pause sites tend to be highly variable, but that their degree of variability is correlated with particular sequence motifs as well as with stress responses. Similarly, steric hindrance of transcription initiation appears to occur at many genes and is intensified in stress responses. Both of these phenomena appear to play major roles in shaping the patterns of aligned reads from NRS data, particularly near the 5′ ends of transcription units.

Our analysis shows, perhaps surprisingly, that even at steady-state it is possible to estimate absolute initiation and pause-escape rates in numbers of events per cell per unit time—provided the average elongation rate at each gene can be estimated or reasonably assumed. Once estimated, these rates can be used, in turn, to obtain various downstream quantities of interest, such as pausing half-lives (**Supplementary Fig. S3**), landing-pad occupanies (**Fig. 4B&C, Supplementary Fig. S12A&B**), or, through combination with other data sources, half-lives of RNA molecules or related quantities [52]. Of course, elongation rates vary across genes and are nontrivial to estimate, but most studies have suggested that they tend to vary by factors of ~2–3, not by orders of magnitude, with average rates fairly close to 2 kb/min in mammalian cells [16, 20, 37]. Therefore, the method appears to be adequate for estimating “ballpark” rates from widely available NRS data even when gene-specific, cell-type-matched elongation rates are not available.

Indeed, our estimates of the half-lives of paused RNAPs—with mean values of 2–3 min. in untreated K562 cells and ~6 min. after heat shock or celastrol treatment (**Supplementary Fig. S3**)—agreed reasonably well with previous experimental estimates. For example, Jonkers et al. measured a mean half-life of 6.9 min. at ~3,200 genes in mouse ES cells [37]; Shao et al. found most half-lives were between 5 and 20 min. at 2,300 genes in *Drosophila* Kc167 cells [22]; and Gressel et al. estimated a median pause duration of 1 min. (corresponding to a half-life of 0.7 min.) for 2,135 genes in CDK9-inhibited human Raji B cells [20]. (The same group subsequently estimated a similar rate for 6,355 protein-coding genes in K562 cells [21].) In addition, Henriques et al. reported half-lives of promoter Pol II complexes from *Drosophila* S2 cells of ~2–15 min at selected loci [53]. However, differences in species, cell types, treatments, and sets of genes make these estimates difficult to compare precisely. Notably, the largest estimates [22, 37] were obtained after treatment with triptolide, which could potentially lead to some inflation in the estimates [26]. Despite these differences, our method seems to be in general agreement overall with previous findings that paused Pol II is typically stable for minutes, and sometimes for tens of minutes, although we detect fewer extremely long half-lives than some previous studies. It is likely that our estimator for *β* reaches saturation when pause peaks become unusually elevated, leading to reduced sensitivity for the extreme tail of the distribution of pausing half-lives. At the same time, our estimator may be more sensitive to short half-lives than some experimental methods.

Despite such limitations, the use of steady-state rather than time-course data also has some important advantages. This approach allows us to analyze all expressed genes, not just a subset at which expression can be induced. In addition, it requires no chemical treatment to block initiation or pause escape, and therefore avoids potential off-target effects. At the same time, it is worth noting that our framework can be extended to the nonequilibrium setting and the use of time-course data [34]. In that setting, it could be used to infer elongation rates together with the other quantities. This version of the model, however, is considerably more complicated mathematically, more difficult to fit to data, and harder to interpret. Additional work will be required before it can be applied to real data.

Our estimator for the pause-escape rate *β* is closely related to the quantity known as the “pausing index” (*I_P_*), which is frequently used to measure the prominence of promoter-proximal pausing [14,35]. In the case where the pause site is constant across cells, we have shown that *I_P_* is precisely the inverse of the maximum likelihood estimator for the rate parameter *β*. When the pause site varies across cells, however, it becomes less straightforward to interpret *I_P_* in physical terms, and a naive interpretation may lead to a strongly biased characterization of the pause-escape rate (e.g., **Fig. 3D**). The details of how *I_P_* is calculated—e.g., by averaging the read counts in the pause region or using their maximum value—also become important in this case. We have shown that our model has a natural extension to variable pause sites, which permits estimation of both the rate parameter *β* and the distribution of pause sites per gene by expectation maximization. In this setting, *β* no longer has a simple relationship to *I_P_*. Interestingly, however, it is possible to find an approximate closed-form relationship in the case where *I_P_* is calculated by averaging, namely, 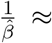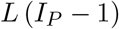, where *L* = *k*_max_ − *k*_min_ + 1 is the length of the pause region (see **Methods**, equation 17). We anticipate that this relationship may help to standardize definitions of *I_P_* and clarify its physical meaning.

Our model also has implications for how to interpret the average read depth in the gene body (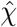 in our notation), which is a natural measure of the transcriptional output at each gene. The model makes clear—as other investigators have previously argued [20]—that 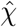 is a measure of the rate of “effective” or “productive” transcription initiation at steady state, rather than of the “potential” rate. That is, in our notation, *χ* is proportional to *ω* rather than *α*. Moreover, in the presence of steric hindrance, *ω* ≈ *ϕβ* (equation 19; see also equation 25), which means that *ω* (and hence *χ*) is effectively limited by the pause-escape rate *β*. Notably, this equation provides a simple and interpretable characterization of the “pause-initiation limit” described by Gressel et al. [20, 21]. It says that the effective initiation rate *ω* can never be greater than the pause-escape rate *β*, with equality when the landing-pad is fully occupied (*ϕ* = 1). The physical meaning of the “potential” initiation rate *α* is somewhat less clear but, at least when *ϕ* is not too close to one, our model does allow extrapolation to a larger rate at which initiation events would hypothetically occur in the absence of steric hindrance. These values can be contrasted with the effective initiation rate in assessing the importance of steric hindrance.

We should emphasize that our model does depend on some crucial simplifying assumptions. In particular, our continuous-time Markov model for the movement of RNAPs along the assumes that any RNAP that successfully initiates will eventually make its way along the entire DNA template. The model excludes the possibility of premature termination, largely because modeling this process—unless its position-dependent rate could somehow be pre-estimated—would make the other rates nonidentifiable. Numerous investigators have argued that pausing dominates in the dynamics of RNAP occupancy, particularly near the promoter, because, among other reasons, promoter-associated RNAP complexes are quite stable and predominantly localize to the pause region [22, 37, 53] (reviewed in [15]). Nevertheless, others have argued, e.g., based on imaging or footprinting techniques, that premature termination may play a much more prominent role [26, 54]. If premature termination does occurs at high rates in the pause region, it could potentially lead to under-estimates of *β* in our framework (because the model will erroneous attributing 5′ peaks completely to pausing), whereas if it occurs at high rates farther along in the gene body, it could lead to under-estimates of both *β* and *χ* (and therefore *ω* and *α*). At the same time, if premature initiation predominantly occurs very quickly and close to the TSS, transiently engaged RNAPs may be largely missed by NGS assays and therefore invisible to our framework, allowing our estimates of *β*, *χ*, and *ω* (but not *α*) to remain accurate. Unfortunately, our steady-state analysis sheds no additional light on this controversy and these questions remain open.

A second process we ignore here is “bursting” of transcription initiation. There is now substantial evidence that rates of transcription initiation do exhibit considerable temporal variation in actively transcribed genes, with apparent oscillations between active and inactive states [23–25, 27, 55, 56]. This pattern is strictly inconsistent with our assumption of a time-homogeneous Markov process. It is worth bearing in mind, however, that our model effectively averages over the processes that are occuring in a large population of cells. Unless those cells are synchronized in their bursting behavior, it would seem that averaging over “on” and “off” states in a bursting model, as our Markov model will effectively do, may be adequate for our purposes—although it is true that bursting could contribute to additional overdispersion in our read-count data.

It is now widely accepted that promoter-proximal pausing is rate-limiting at many genes and this step is frequently regulated [15]. Our model helps to make clear, however, that steric hindrance can modulate this important regulatory step in critical ways. On one hand, the model confirms that, in the presence of rate-limiting pause-escape (small *β*), promoting initiation (increasing *α*) will have little or no impact on transcriptional output, because the pause-initiation limit will quickly be reached. In this setting, a more rapid and effective transcriptional response will come from releasing RNAPs from the paused state (increasing *β*). At the same time, the model also reveals that the dynamic equilibrium between initiation and pause-escape will tend to limit the *duration* of the transcriptional response to pause-release. The reason is that the geometry of the pause region allows only a few RNAPs to be kept “in the chamber,” ready to be released into productive elongation. As soon as these ~1–3 RNAPs are released in each cell, initiation will again become rate-limiting. Thus, the effects of changes to initiation and pause-release cannot be fully separated; rather, the two processes interact with one another in a kind of dance, each limiting the other under certain circumstances. For this reason, each process is the natural target for regulation of gene expression in a different regime—for example, initiation when a sustained transcriptional response is required, and pause-escape when a rapid and/or synchronous but transient response is needed. Rates of initiation and pause-escape are perhaps best thought of as two “knobs” for control of a single interrelated dynamical system near the TSS of each gene.

## Methods

### Continuous-time Markov Model

The probabilistic model (**Fig. 1A&B**) consists of a continuous-time Markov model that describes the stochastic movement of individual RNAPs along the DNA template, and a conditional generating process by which read counts arise independently at each site in proportion to RNAP occupancy (defined below). The Markov model consists of *N* + 1 states corresponding to the *N* possible nucleotide positions of the active site of an RNAP as it moves along an *N*-nucleotide DNA template, plus an additional state (labeled 0) that abstractly represents “free” RNAPs, not currently engaged in transcription and available for new initiation events. Each state *i* in the Markov model corresponds to a binary random variable *Z_i_*, indicating whether the RNAP is (*Z_i_* = 1) or is not (*Z_i_* = 0) at position *i* at a particular time *t* (**Fig. 1B**). In our setting, the use of this time-homogeneous model depends on two key assumptions: (1) that collisions between RNAPs are rare, allowing the movement of each RNAP to be considered independently of the others; and (2) that premature termination of transcription is sufficiently rare that each RNAP can be assumed to traverse the entire DNA template if it is given enough time (see Discussion for limitations).

The model distinguishes between two segments of each transcription unit: (1) the first *k* nucleotides, known as the *pause peak*, where RNAP tends to accumulate owing to promoter-proximal pausing (typically *k* ≈ 50) [9]; and (2) the subsequent *N* − *k* nucleotides, where RNAP tends to be relatively unimpeded, which is typically referred to as the *gene body*. Movement of the RNAP is defined by four rate parameters: an initiation rate *α* (from state 0 to state 1), a pause-escape rate *β* (from state *k* to state *k* + 1, a termination rate *γ* (from state *N* to state 0), and a constant per-nucleotide elongation rate *ζ* (for all other allowable transitions). Because the states must be visited in a sequence, the infinitesimal generator matrix for the Markov chain **Q**= {*q_ij_*} has a simple form, with positive terms only on the diagonal *q_i,j_* such that *j* = *i*+1, negative terms on the main diagonal, and zeroes elsewhere. For mathematical convenience, we assume that the initiation, pause-escape, and termination steps are coupled with single-nucleotide elongation steps and occur at rates *ζα*, *ζβ*, and *ζγ*, respectively. As a result, as long as *ζ* is the same across nucleotides, it can be considered a scaling factor that applies equally to all steps in the process. Specifically,

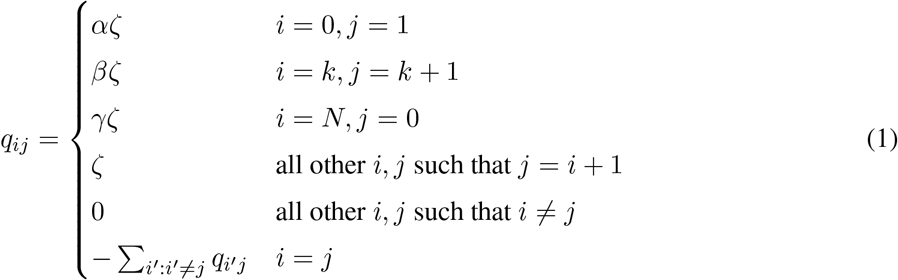

where element *q_ij_* indicates the instantaneous rate at which an RNAP transitions from state *i* to state *j*, and by convention, the values along the main diagonal are set such that the rows sum to zero.

### Stationary Distribution

This continuous-time Markov model allows for calculation of the probability of transitioning from any state *i* to any other state *j* over a period of time *t* ≥ 0 (see [34]), but in this article we are interested in steady-state conditions and therefore focus on the limiting distribution for ending states as *t* → ∞. This stationary distribution, denoted ***π***, is invariant to *ζ* and can be found easily by solving the equation ***π*Q= 0**. Because state 0 is simply an abstraction to allow for the recirculation of RNAP, we omit it and describe the stationary distribution conditional on RNAP occupancy along the DNA template. This conditional stationary distribution can be expressed as ***π*** = (*π*_1_*,…, π_N_*) such that,

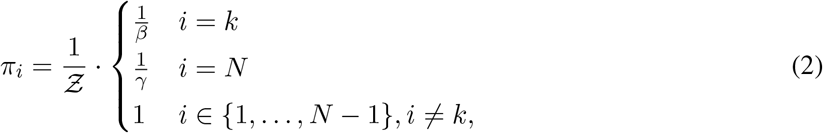

with normalization constant 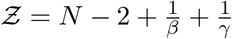.

This distribution has an intuitive interpretation. First, it is natural that, conditional on RNAP occupancy, the steady-state distribution is invariant to both *α* (which defines the rate at which occupancy is initiated but has no effect thereafter) and *ζ* (which defines the “flow” along the DNA template but does not favor one nucleotide position over another). In addition, as a result of local slowdowns in elongation, *π_i_* is elevated relative to the gene body by factors of 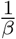 and 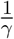 at the pause peak and termination peak, respectively. Notice that both peaks take the form of “spikes” at single nucleotide positions under this model; in later sections we will generalize the model to allow for a broader pause peak.

When comparing different transcription units, we have to allow for differences in TU-specific initiation and elongation rates. In particular, in addition to obeying equation 2, the relative RNAP densities at TU *j* will be proportional to 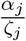, where *α_j_* and *ζ_j_* are the TU’s initiation and elongation rates, respectively. Furthermore, as detailed below, estimation of these rates is confounded by the sequencing depth, *λ*. Because these parameters are not identifiable at steady state, we represent them by the compound parameter 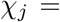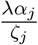.

In addition, the termination peak tends to be difficult to characterize with real nascent RNA sequencing data, owing to transcriptional run-on, poorly characterized 3′ ends of genes, and other factors. Therefore, from this point on, we omit the parameter *γ* and assume *N* is defined such that ambiguities at the 3′ ends of TUs are excluded.

With these assumptions, when comparing the RNAP densities at various sites *i* along various TUs *j*, we expect them to be proportional to,

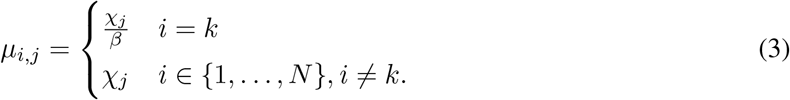

### Generative Model for Sequence Data

To allow for the fact that the RNAP positions are only indirectly observed through the sequencing process, we add a second layer to the model that describes the probabilistic process by which sequencing read counts are generated conditional on an underlying density of RNAP at each nucleotide, as defined by the continuous-time Markov model. In this way, we obtain a full generative model for the observed sequence data that is defined by the model parameters, enabling inference of all parameters from the data.

Because we have freedom in how to set the read-depth scaling parameter *λ* (see below), we simply take *μ_ij_* (as defined in equation 3) to be the expected read depth at position *i* of TU *j*. We then assume that the read count *X_i,j_* is Poisson-distributed with this mean. It is possible to use other generating distributions to allow for overdispersion of the read counts [34], but the Poisson assumption seems to be adequate in our case and it is particularly convenient for parameter inference.

With these assumptions, let the data for a single TU be denoted **X**= (*X*_1_*,…, X_N_*), where *X_i_* represents the number of sequencing reads having their 3′ end aligned to position *i*. (We omit the *j* index in this case to simplify the notation.) Assuming conditionally independent Poisson distributions at each site, the steady-state log likelihood function for a single TU is given by,

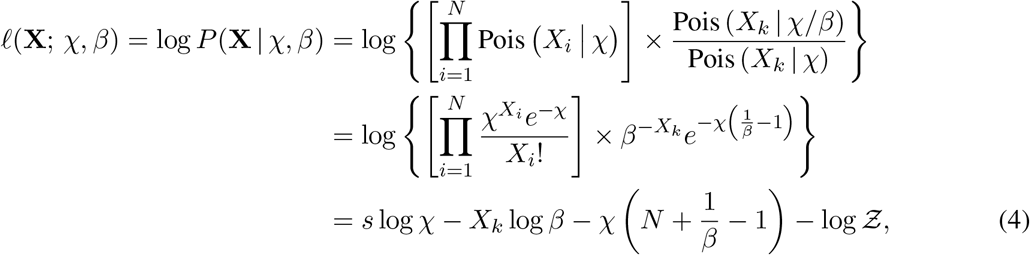

where 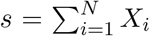 is a statistic equal to the sum of all read counts and 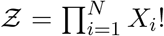 is a normalization term that does not depend on the model parameters and can be ignored during optimization.

### Inference at Steady-State

Under this likelihood function (equation 4), the maximum-likelihood estimators for *χ* and *β* have simple closed-form solutions:

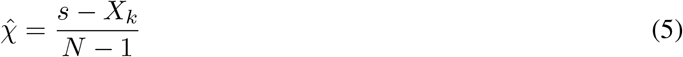

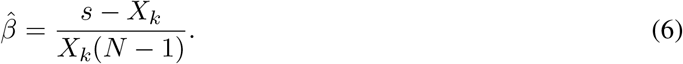

Notice that the estimator for *χ* is simply the average read depth, excluding the pause peak, and the estimator for *β* is the ratio of that same average read depth to the read depth in the pause peak. These are estimators that have been widely used in the analysis of nascent RNA sequencing data, with more heuristic justifications.

In practice, it tends to be better to avoid the complex signal in the pause region in estimating *χ* and instead estimate it from a downstream portion of the gene body. We define 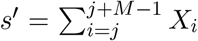, where *M* is the length of the interval considered, and estimate *χ* and *β* as,

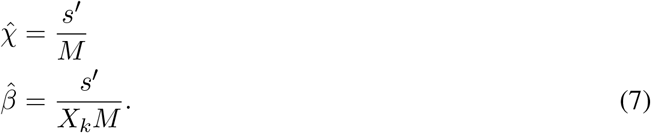

For the remainder of the paper, we assume the use of these simpler, more robust estimators for *χ* and *β*.

### Allowing for Variation in the Pause Site

In real samples, the pause site tends to vary across cells, leading to a broad pause peak in nascent RNA sequencing data. We address this complication by allowing the location of the pause peak, *k*, to vary between a *k*_min_ and a *k*_max_ according to an appropriate distribution, and then assuming each read count *X_k_* within this range reflects a mixture of cells that do and do not have their pause site at position *k*. Assume that, for *k* ∈ {*k*_min_*,…, k*_max_}, *f_k_* represents the fraction of cells with pause site *k*, and that the read count *X_k_* derives from a mixture of one Poisson distribution with rate *χ/β* · *f_k_* and a second Poisson distribution with rate *χ* · (1 − *f_k_*). If we denote by *Y_k_* the (unknown) portion of the read count that derives from the first process, then the log likelihood function can be expressed as (cf. equation 4),

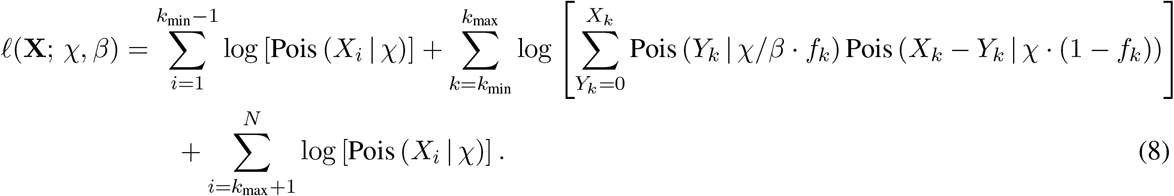

This log likelihood function does not have a closed-form solution but it is straightforward to maximize by expectation maximization. The complete-data log likelihood function, with known values of *Y_k_*, can be expressed in terms of compact sufficient statistics (analogous to equation 4) as,

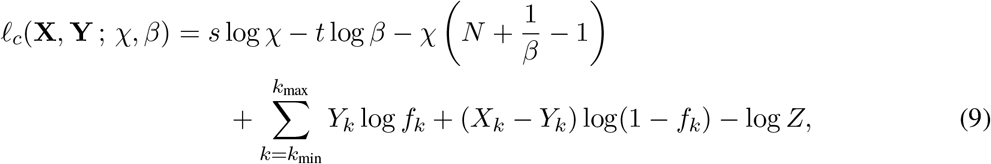

where *s* = Σ_*i*_ *X_i_* and *t* = Σ_*k*_ *Y_k_*. The expected value of this function, averaging over the latent variables *Y_k_*—the quantity to maximimize in EM—is simply,

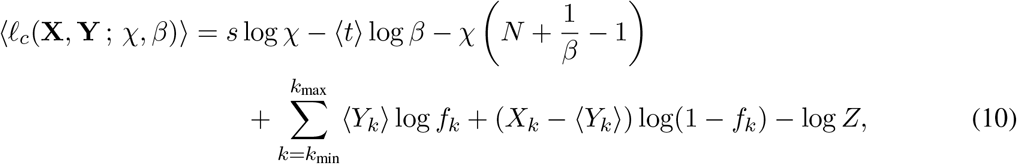

where 〈*Y_k_*〉 and 〈*t*〉 = Σ_*k*_ 〈*Y_k_*〉 denote the posterior expected values of *Y_k_* and *t*, respectively. For simplicity, assume that *χ* is pre-estimated for a portion of the gene body downstream of *k*_max_ using equation 7. If in addition the values of *f_k_* are fixed, then *β* can be simply estimated as,

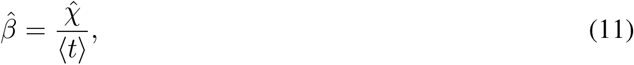

The values 〈*Y_k_*〉 can be computed by observing that, because *X_k_* is the sum of two Poisson-distributed variables, *Y_k_* | *X_k_* is binomially distributed with probability,

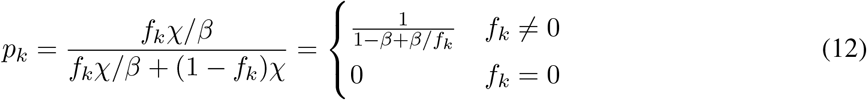

Therefore, 〈*Y_k_*〉 = *X_k_* · *p_k_*, and

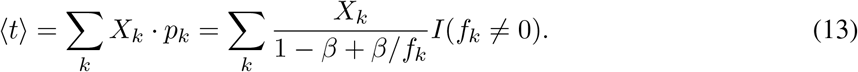

Thus, an EM algorithm can be implemented by iteratively applying equations 11 and 13, in turn, until convergence.

Furthermore, to estimate the distribution of pause sites from the data at each TU, we assume the *f_k_* values have a truncated Gaussian distribution with mean *μ* and variance *σ*^2^,

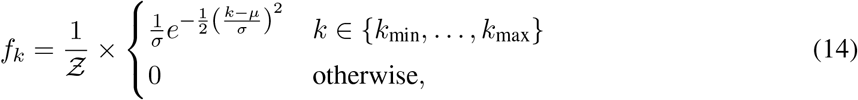

where the explicit normalization constant Z is needed because the distribution is applied to a bounded interval and is defined at integer values only.

In this case, the EM updates for *μ* and *σ*^2^ are simply,

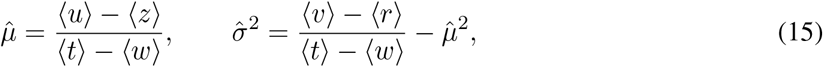

where,

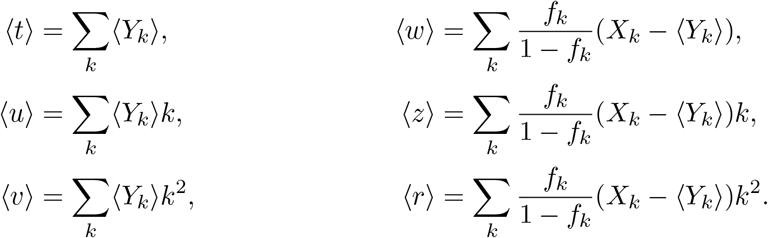

Notice that all of these quantities are easily computed from the 〈*Y_k_*〉 values together with *f_k_* and *X_k_*.

### Approximate Relationship between *β* and the Pausing Index *I_P_*

The model above requires iterative estimation, but an approximate closed-form expression for *β* in terms of the pausing index *I_P_* can be obtained by noting that the excess reads in the pause region, *t*, can be estimated reasonably well by multiplying the difference between the average read depths in the pause region and gene body by the length of the pause region, *L* = *k*_max_ − *k*_min_ + 1:

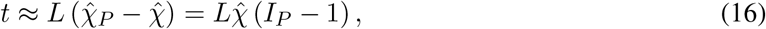

where 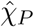 is the average read depth in the pause region and 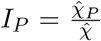 is the pausing index as computed by averaging across the pause region. Therefore,

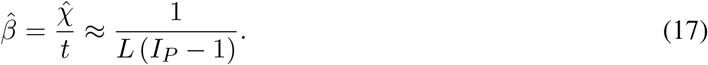

Thus, an interpretable pause-escape rate parameter can be estimated approximately from the pausing index *I_P_*. This equation provides a physical interpretation for *I_P_* and also explains why our naive initial estimation strategy tended to over-estimate *β* by a factor of approximately *L*. This strategy, however, does not allow estimation of the mean or variance of the pause site at each gene.

### Allowing for Steric Hindrance of Initiation

To accommodate steric hindrance of initiation at steady state, we introduce a distinction between a *potential* rate of initiation in the absence of occlusion of the initiation site, *α*, and the *effective* rate of initiation after a portion of initiation events are blocked by an existing RNAP molecule, which we denote *ω*. We assume *ω* = (1 − *ϕ*)*α*, where *ϕ* is the probability of that the “landing pad” required for a new initiation event is already occupied by an RNAP. Thus, *ω* ≤ *α*. Notice that any estimation of initiation rates based on the density of RNAPs in the gene-body will be representative of *ω*, not *α*; a correction may be required to estimate *α* accurately.

We first assume that the “footprint” of an engaged polymerase, *ℓ*, is sufficiently large that at most one RNAP can be present in this region at a time. (We relax this assumption below.) We further assume that elongation up to position *k* occurs much faster than the initiation rate *αζ* or the pause-escape rate *βζ*, so that the dynamics of elongation through the pause peak can be ignored. In this case, occupancy of the landing pad can be described using a simple two-state continuous-time Markov model (**Fig. 5A**). Here, either the landing pad is unoccupied (state 0) and is therefore available for new initiation events, which occur at rate *αζ*; or the landing pad is already occupied by an RNAP (state 1) and no new initiation events can occur until that RNAP escapes from the pause site, which occurs at rate *βζ*. Thus, at steady state, the landing-pad occupancy *ϕ* is simply given by the stationary distribution of the occupied state,

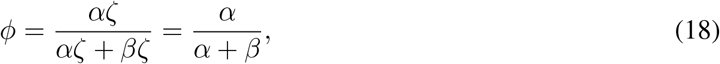

and the effective initiation rate, allowing for steric hindrance, is given by,

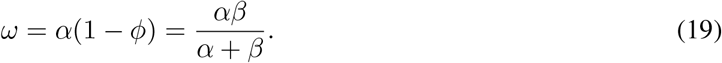

**Figure 5:**
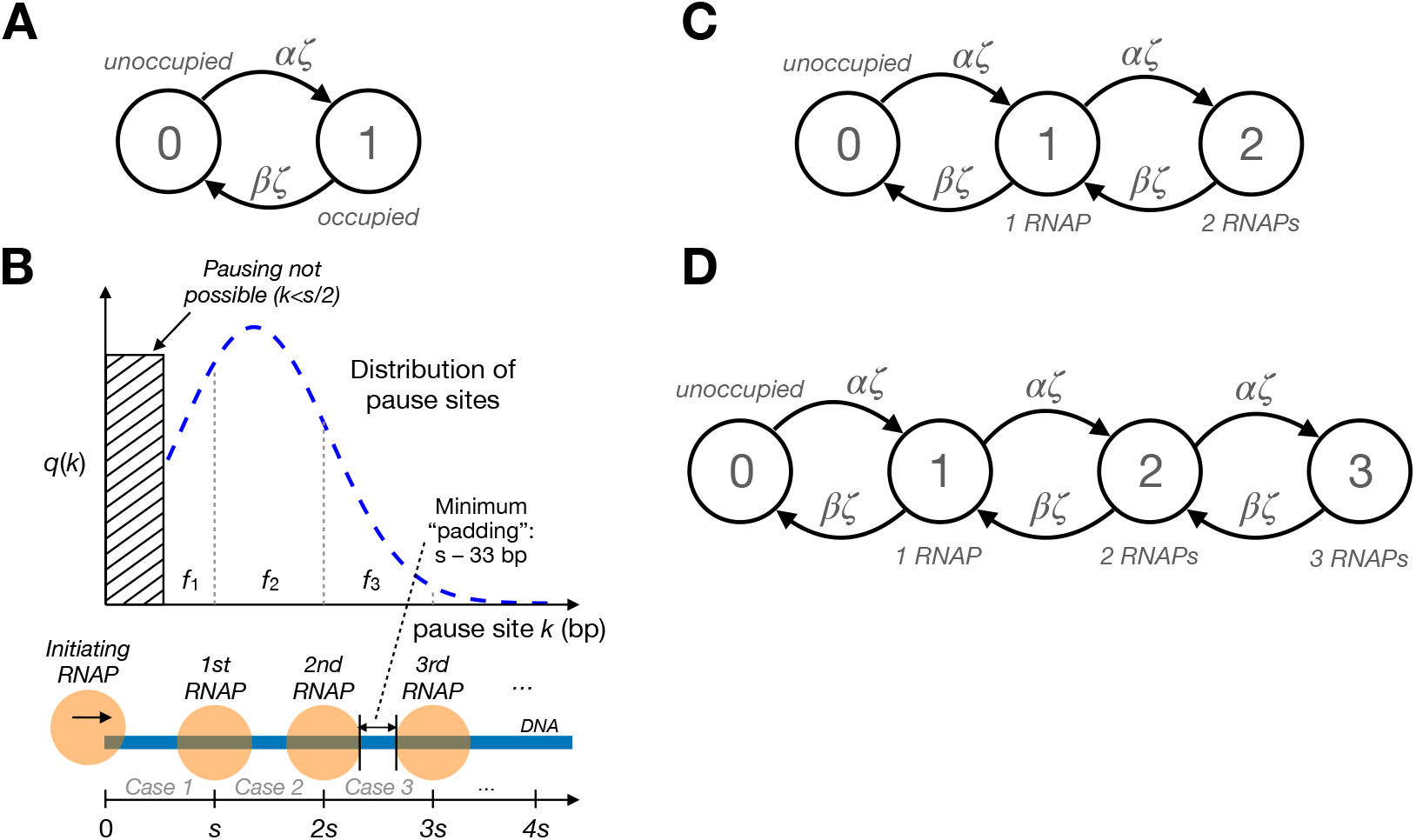
**A**. Two-state continuous-time Markov model for steric hindrance of transcriptional initiation, assuming at most one RNAP at a time in the pause region. The pause region must be either unoccupied (state 0) or already occupied by another RNAP (state 1). Transitions from state 0 to state 1 occur at the (unimpeded) initiation rate, *αζ*, and transitions from state 1 to state 0 occur at the pause-escape rate, *βζ*. The stationary frequency of state 1 defines the landing-pad occupancy *ϕ* and is given by 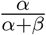. **B**. Illustration showing a hypothetical distribution of pause sites *k* and its implications for the number of RNAPs that can simultaneously occupy the pause region. When *k* ≤ *s*, where *s* is the minimum center-to-center spacing between adjacent RNAPs, only one RNAP is possible (Case 1 in the text); when *s < k* ≤ 2*s*, up to two are possible (Case 2); and when 2*s < k* ≤ 3*s*, up to three are possible (Case 3). Notice that the portion of the density corresponding to each Case *r* is given by *f_r_*. **C**. Generalization of Markov model to accommodate up to two RNAPs in the pause region (Case 2). **D**. Further generalization to accommodate up to three RNAPs (Case 3). The equation for *ϕ* can be generalized to account for these cases (see text).

Notice that these equations also imply that *ω* = *ϕβ*, meaning that, at steady state, the effective initiation rate *ω* must always be less than or equal to the pause-escape rate *β*, with *ω* approaching *β* as the landing-pad occupancy *ϕ* approaches unity. Therefore, if one estimates an effective initiation rate 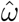 and a pause-escape rate 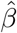 from the data, as described above, then one can obtain estimates of *ϕ* and *α* as follows,

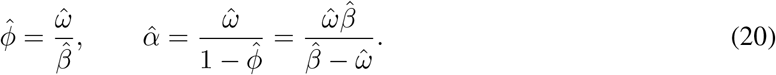

Notice that the estimator for *ϕ* will be proportional to the read-counts in the pause peak (see, e.g., equation 7).

### Steric Hindrance with Multiple RNAPs

As it turns out, the assumption of ≤1 RNAPs per pause region is often too restrictive, and the presence of more than one RNAP in this region can have a substantial impact on the landing-pad occupancy *ϕ*. In this section, we generalize the model for steric hindrance to allow for any number *r* of RNAPs in the pause region, focusing in particular on the case of *r* ≤ 3, which we expect to cover essentially all plausible scenarios in human cells.

Let *s* be the minimum center-to-center spacing, in nucleotides, between adjacent RNAPs on the DNA template. As noted in the main text, structural data suggests *s* is at least 33 bp but more plausibly *s* ≈ 50 bp [18, 19, 41, 46]; we also consider the case of *s* = 70 bp for comparison. We further assume that a new initiation event can successfully occur if, and only if, the previous RNAP has advanced to a position *i > s* (in other words, the “landing pad” for new initiation events has size *£* = *s*). Consequently, the maximum possible number of RNAPs in the pause region is *r* = 1 if the pause site *k* ≤ *s*, *r* = 2 if *s < k* ≤ 2*s*, *r* = 3 if 2*s < k* ≤ 3*s*, and so on (see **Fig. 5B**). In general, 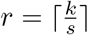.

In addition, we assume a probability mass function *q*(*k*) for the pause site *k* across cells, with cumulative distribution function 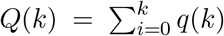. Let *f*_1_ be the density associated with *r* = 1; that is, *f*_1_ = *Q*(*k* = *s*). Similarly, *f*_2_ is the density associated with *r* = 2, *f*_2_ = *Q*(*k* = 2*s*) − *Q*(*k* = *s*); and *f*_3_ = *Q*(*k* = 3*s*) − *Q*(*k* = 2*s*). In general, *f_r_* = *Q*(*k* = *rs*) − *Q*(*k* = (*r* − 1)*s*) (**Fig. 5B**).

Now, in each cell, *k* has a single value and therefore one of a series of mutually exclusive cases must apply. Let us denote by Case *r* (for *r* ∈ {1, 2, 3,…}) that the maximum possible number of RNAPs is *r*. Thus, in Case 1, *r* = 1 and *k* ≤ *s*; in Case 2, *r* = 2 and *s < k* ≤ 2*s*; and so on. Given *q*(*k*), we know that Case *r* occurs with probability *f_r_*. Therefore, we can calculate *ϕ* as a mixture of case-specific landing-pad probabilities, 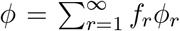, where *ϕ_r_* is the probability that the landing pad (first *s* nucleotides) is occupied in case *r*. In practice, we approximate this quantity as 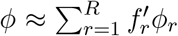 where *R* is the maximum plausible value of *r* (here, *R* = 3) and,

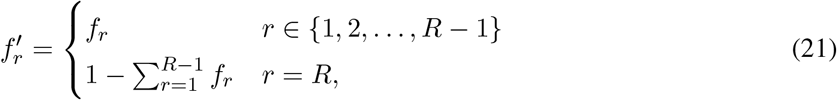

where the last term, *f′_R_* is “rounded up” to account for the remaining tail of the distribution.

The case of *r* = 1 has already been described in the previous section. It can be captured by a two-state model, where the landing-pad is either unoccupied (state 0) or occupied by a single RNAP (state 1; **Fig. 5A**). Therefore,

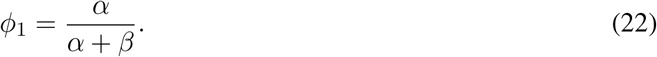

It turns out that the cases of larger values of *r* follow naturally through the addition of more states. For example, when *r* = 2, the landing pad is either unoccupied (state 0), occupied by one RNAP (state 1), or occupied by two RNAPs (state 2). Assuming no two events can occur simultaneously, state 2 can be reached only when a new initiation event occurs while state 1 is occupied (at rate *αζ*), and a pause-release event in state 2 causes a return to state 1 (at rate *βζ*). The result is a chain of states as shown in **Fig. 5C**. In addition, under the assumption that elongation through the pause peak is instantaneous, the landing pad is occupied if, and only if, state 2 is occupied. Thus, *ϕ*_2_ is given by the stationary frequency of state 2 in this model, which can be shown to be,

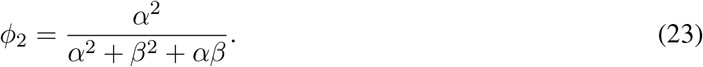

Similarly, the case of *r* = 3 can be addressed by extending the chain further with a state 3, and setting *ϕ*_3_ equal to the stationary frequency of that state (**Fig. 5D**),

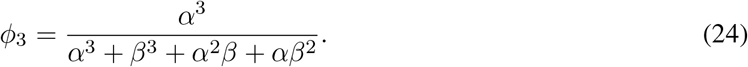

Therefore, assuming *R* = 3, we can estimate *ϕ* as,

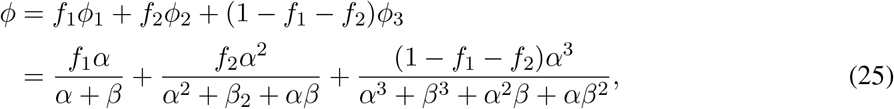

allowing for the possibilities of one, two, or three RNAPs in the pause region of each cell. As *s* grows larger and/or *q*(*k*) shifts toward the TSS, the fraction *f*_1_ approaches 1, and equation 25 approaches equation 18.

As it turns out, substitution of *ω*/(1 − *ϕ*) in place of *α* in this more general expression for *ϕ* leads to a complex polynomial that cannot be easily solved for *ω*. Instead, we solve this equation numerically to obtain *ϕ* and *α* from *ω* and *β* (analogous to equation 20).

### Fitting the Steric-Hindrance Model to Data

The multi-RNAP steric hindrance model can be combined with the model for variable pause sites across cells and fitted to the data by a relatively straightforward extension of the EM algorithm described above. As in that case, we assume that the unscaled initiation rate, *χ*, is pre-estimated from data in the gene body. In addition, however, this case requires pre-estimation of the scale-factor *λ* and the elongation rate *ζ*, so that a scaled estimate of the effective initiation rate, 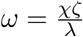, can be obtained from each estimate of *χ*. In practice, we obtain these scale factors by calibrating with respect to other studies (see“Calibrating the initiation rate,” below).

In addition, we must address the problem that the landing-pad occupancy *ϕ* is constrained to fall between 0 and 1, but the relationships above do not enforce such a constraint. For example, in the single-RNAP case, where 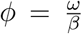 (equation 20), *ϕ* will be undefined whenever *ω > β*. A similar (but more complicated) relationship holds in the multi-RNAP case.

To address this problem, we take a Bayesian approach and assume a (weakly) informative prior distribution for *ϕ*. This strategy not only restricts *ϕ* to the allowable range but has the benefit of regularizing the model when information in the data about *ϕ* is weak. With the assumption of a Beta(*ϕ* | *a, b*) prior for *ϕ*, with shape parameters *a* and *b* (we assume *a* = *b* = 2 throughout), the likelihood (cf. equation 8) becomes,

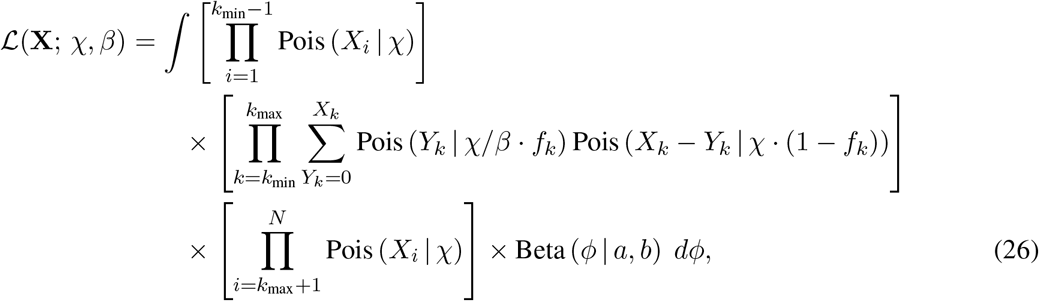

and the expected complete-data log likelihood (cf. equation 10) becomes,

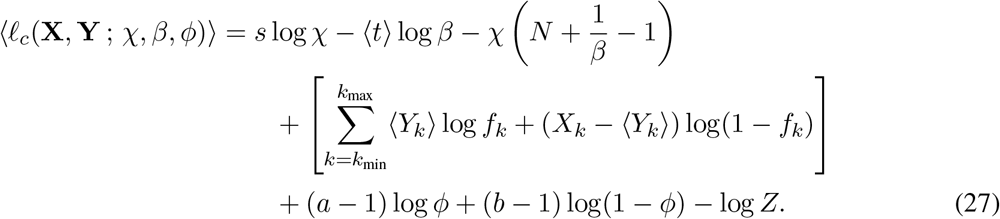

From a comparison of equations 10 and 27, it is evident that the calculation of the summary statistics *〈t〉, 〈u〉, 〈v〉, 〈w〉, 〈z〉*, and *〈r〉* and the updates for *μ* and *σ*^2^ will all remain unchanged (equations 12–15). However, this simplified presentation obscures that the parameters *ϕ* and *β* are implicitly linked by a function, *ϕ* = *g*(*β*; *χ*), which is indirectly defined by equation 25 as well as by the relationship *ω* = (1 − *ϕ*)*α*. Thus, the update for *β* in this case is no longer that shown in equation 11 but now must also consider the terms that depend on *ϕ*. As it turns out, in the full multi-RNAP model, rewriting equation 27 in terms of *β* only (i.e., by substitution for *ϕ*) leads to rather unwieldy polynomial expressions. Nevertheless, the M-step in the EM algorithm can be performed numerically without much trouble. In particular, on each iteration of the algorithm, we calculate all expected sufficient statistics as before (E-step), but then, for the M-step, we estimate *β* by numerically maximizing the portion of equation 27 that depends on *β* and *ϕ*, that is,

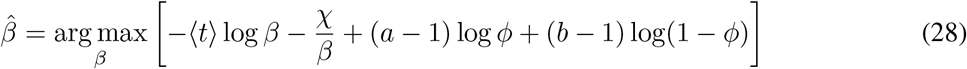

subject to the constraints of equation 25 and the relationship *ω* = (1 − *ϕ*)*α*. Thus, the previous EM algorithm can be adapted to the multi-RNAP steric hindrance case by simply replacing the M-step for *β* with this numerical optimization. No other changes are required.

### SimPol Simulator

SimPol (“Simulator of Polymerases”) tracks the independent movement of RNAPs along the DNA templates of a large number of cells. It accepts several key user-specified parameters, including the initiation rate, pause-escape rate, a constant or variable elongation rate, the mean and variance of pause sites across cells, as well as the center-to-center spacing constraint between RNAPs (*s*), the number of cells being simulated, the gene length, and the total time of transcription. The simulator simply allows each RNAP to move forward or not, in time slices of 10^*−*4^ minutes, according to the specified position-specific rate parameters. It assumes that at most one movement of each RNAP can occur per time slice. The simulator monitors for collisions between adjacent RNAPs, prohibiting one RNAP to advance if it is at the boundary of the allowable distance from the next. After running for the specified time, SimPol outputs a file in bigWig format that records all RNAP position. SimPol is written in the statistical programming language R [57], and depends on the optparse [58], Matrix [59], and rtracklayer [60] packages.

### Generation of Synthetic NRS Data

Using SimPol, we simulated genes 2000 bp in length with initiation (*αζ*) and pause-escape (*βζ*) rates that spanned two orders of magnitude, ranging from 0.1 to 10 events per min. per cell. Elongation rates at each nucleotide position were randomly sampled from a truncated normal distribution, with mean = 2,000 bp/min, sd = 1000 bp/min, min = 1500 bp/min and max = 2500 bp/min. When a fixed pause site was assumed, it occurred at position *k* = 50 bp; variable pause sites assumed a truncated normal distribution for *k*, with mean = 50 bp, sd = 25 bp, min = 17 bp and max = 200 bp. Our main simulations assumed a center-to-center spacing *s* = 50 bp, but alternative simulations assumed *s* = 33 bp and *s* = 70 bp. For each parameter combination, we simulated 20,000 cells for the equivalent of 40 min. (400,000 time slices), which appeared to be sufficient to reach equilibrium in all cases.

Based on the output of each SimPol run, we randomly sampled 5,000 of the 20,000 cells. This sampling step was performed 50 times for each run. To save in computation, the same source collection of 20,000 cells was used for each replicate, after veryifying that it made little difference to rerun the full simulation to equilibrium each time. For each of these replicates, we then sampled a read count at each position *i* from a Poisson distribution with mean *μ_i_* such that,

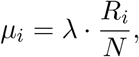

where *R_i_* is the number of sampled cells having an RNAP whose active site (center) is at position *i* and *N* = 5, 000 is the total number of sampled cells. The scale parameter *λ* was calibrated such that a typical choice of initiation (*αζ* = 1 event/min) and pause-escape (*βζ* = 1 event/min) rate parameters resulted in an average read depth of 0.049 reads/bp in the gene body, as we observed in the real data from ref. [17] (median value).

For simplicity, we performed this step separately at each nucleotide only in the pause region (the first 200 nucleotides). Because the RNAP densities throughout the gene bodies are fairly homogeneous, and the corresponding read counts are averaged anyway, we sampled the total read count for each gene body in one final step by scaling the Poisson distribution appropriately. This strategy also allowed us to extrapolate from our 2,000-bp simulated genes to genes of more realistic length. Specifically, the total read count for the gene body was sampled from a Poisson distribution with mean *μ*_GB_ such that,

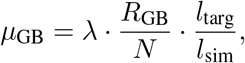

where *R*_GB_ is the total number of RNAPs across the simulated gene body, *N* = 5, 000 is the total number of cells sampled, *l*_targ_ = 19, 800 bp is the target gene-body length, and *l*_sim_ = 1, 800 bp is the simulated length. Notice that the scale parameter *λ* was held fixed throughout so that average read-depths would increase or decrease appropriately as the rate parameters were altered.

### Analysis of Real Data

We obtained published K562 PRO-seq libraries from the heat shock [36] and celastrol studies [17] and processed them using the PROseq2.0 pipeline (https://github.com/Danko-Lab/proseq2.0) in single-end mode [61]. The 3′ ends of reads—which approximately represent the active sites of isolated RNAPs—were recorded in bigWig files and used for analysis. Mapping was performed with human genome assembly GRCh38.p13 and gene annotation were downloaded from Ensembl (release 99) in GTF [62]. Annotations of protein-coding genes from the autosomes and sex chromosomes were used, excluding overlapping genes on the same strand. To improve TSS positioning, we augmented the gene annotations with CoPRO-cap (Coordinated Precision Run-On and sequencing with 5′ Capped RNA) data for K562 cells from ref. [44] (see [21, 63] for similar uses of GRO-cap data). In particular, we used the position with highest CoPRO-cap signal within a 250 bp radius around each annotated TSS as a refined TSS and discarded genes for which no CoPRO-cap signal was found. We then considered the 200 bp starting at this refined TSS as the “pause region,” and the region from 1,250 bp to up to 90 kbp downstream (but not past the annotated end of the gene) of this TSS as the “gene body.” We excluded any gene with fewer than 20 reads mapped to either the pause peak or the gene body. For the remaining genes, the read counts at each of 200 positions in the pause region, and the total read counts in the gene body, were summarized in a table and used for all downstream analyses.

### Calculation of half-lives

RNAP half-lives were calculated from estimates of the pause-escape rate *β* and average values of the pause-site position *k* by the following equation,

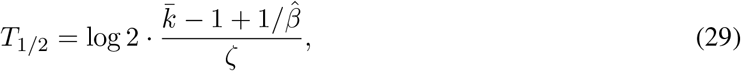

where 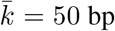 *ζ* = 2000 bp/min, and 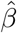 is the estimated pause-escape rate. Here, the quantity 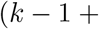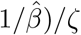 represents the expected time required for an RNAP to pass through the pause region and the pause site, and the factor log 2 converts the mean of an exponential distribution to a half-life.

### Discriminative motif finding

DNA sequences 200 bp downstream of the TSS were extracted, and STREME [64] was used to identify the motifs enriched in the 10% genes with narrowest pause peaks (smallest 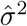) compared with the 10% of genes with broadest peaks (largest 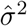). Tomtom [65] was then used to annotate the results with known motifs from HOCOMOCO v11 [66].

To validate the sequence motif enrichment of TAF1, ChIP-seq signals for TAF1 in K562 cells downstream of the TSS were extracted separately for genes exhibiting narrow and broad peaks. These signals were plotted using the R package Genomation [67]. Sequence logos for regions around the position with maximum PRO-seq read counts in the pause peak were plotted using the R package ggseqlogo [68].

### Calibrating the initiation rate

As described in the text, we made use of a “low” (L) calibration of 0.2 initiation events per minute [47–50], and a “high” (H) calibration based on ref. [20], who reported a median of 1.7 initiation events per minute for mRNAs. These calibrations were performed separately for each of our four analyzed data sets: the untreated and treated samples from the heat-shock [36] and celastrol [17] studies. We performed all calibrations using “housekeeping” (HK) genes from ref. [51], to minimize sensitivity to differences between cell types and other properties. We first identified a subset of genes that fell in the intersection of the HK set, the genes from ref. [20], and each of our sets. This subset numbered between 443 and 446 genes for our four data sets. In each case, for the L calibration, we simply scaled our *ω* values so that the median value of *ωζ* within this set was equal to 0.2 events/min. For the H calibration, we scaled our *ω* values so that the median value of *ωζ* within this set matched the median value reported by Gressel et al. [20] for the same subset of genes, which ranged from 2.36–2.37 events/min for our four data sets. Finally, to focus on genes exhibiting robust expression in both the treated and untreated samples, we identified a subset of genes that fell in the top 80% by 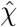 in both samples. This filter resulted in a total of 6,087 genes for the heat-shock data set and 5,964 genes for the celastrol dataset.

## Data Availability

PRO-seq fastq files of the heat shock [36] and celastrol [17] datasets were downloaded from GEO with accession numbers GSE89230 and GSE96869. BigWig files of CoPRO-cap were downloaded with accession number GSE116472 [44]. The bigWig file for TAF1 Chip-seq was downloaded from ENCODE with accession number ENCFF101GBL [69]. All data are for human K562 cells, and replicates were combined for analysis.

## Acknowledgments

We thank Charles Danko, Gilad Barshad, and other members of the Siepel and Danko laboratories for helpful discussions. We also thank Michael Lidschreiber and Patrick Cramer for providing raw data from refs. [20] & [21]. This research was supported by US National Institutes of Health grants R35-GM127070 and R01-HG009309 (to AS), and by the Simons Center for Quantitative Biology at Cold Spring Harbor Laboratory. The content is solely the responsibility of the authors and does not necessarily represent the official views of the US National Institutes of Health.

## Notes

### Competing Interest Statement

The authors have declared no competing interest.

