## Supplementary Information for "Model-based characterization of the equilibrium dynamics of transcription initiation and promoter-proximal pausing in human cells"

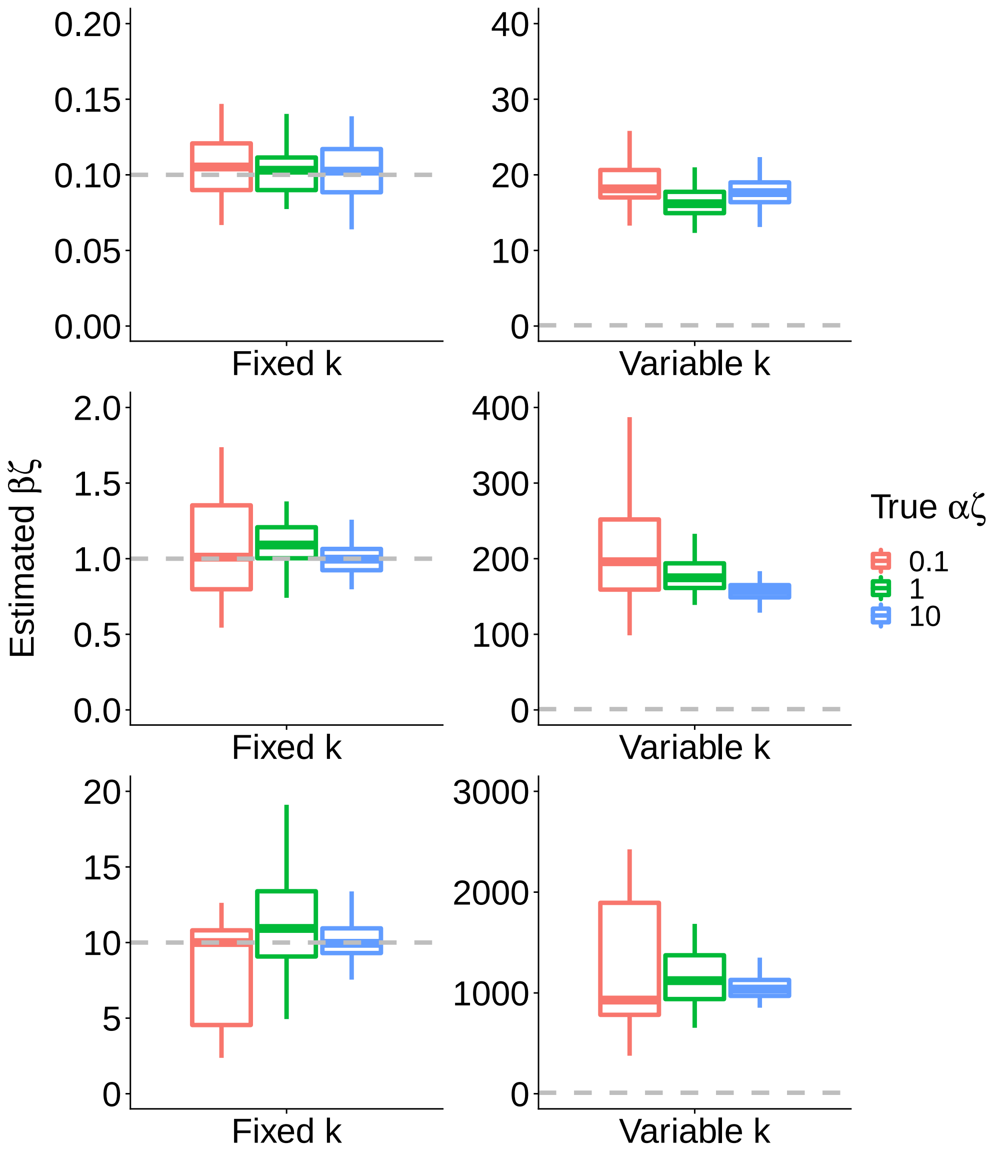


Supplementary Figure S1: Estimated values of $\beta\zeta$ under the initial model for simulated true values of $\beta\zeta \in \{0.1, 1.0, 10\}$ (top to bottom) and $\alpha\zeta \in \{0.1, 1, 10\}$ (see key), when the pause-site $k$ is fixed (left) or variable across cells (right) in simulation. This is an expanded version of **Fig. 2B** showing a broader range of $\beta\zeta$ values. As in that case, dashed lines indicate the ground truth; boxplots summarize 50 replicates of the simulation; box boundaries indicate 1st and 3rd quartiles, and horizontal line indicates median. A value of $\zeta = 2$ kb/min is assumed so that $\alpha\zeta$ and $\beta\zeta$ can be assumed to have units of events per minute. Pause sites occur at a mean position of $k = 50$ bp. In the variable case, we assume a Gaussian distribution with a standard deviation of 25 bp.

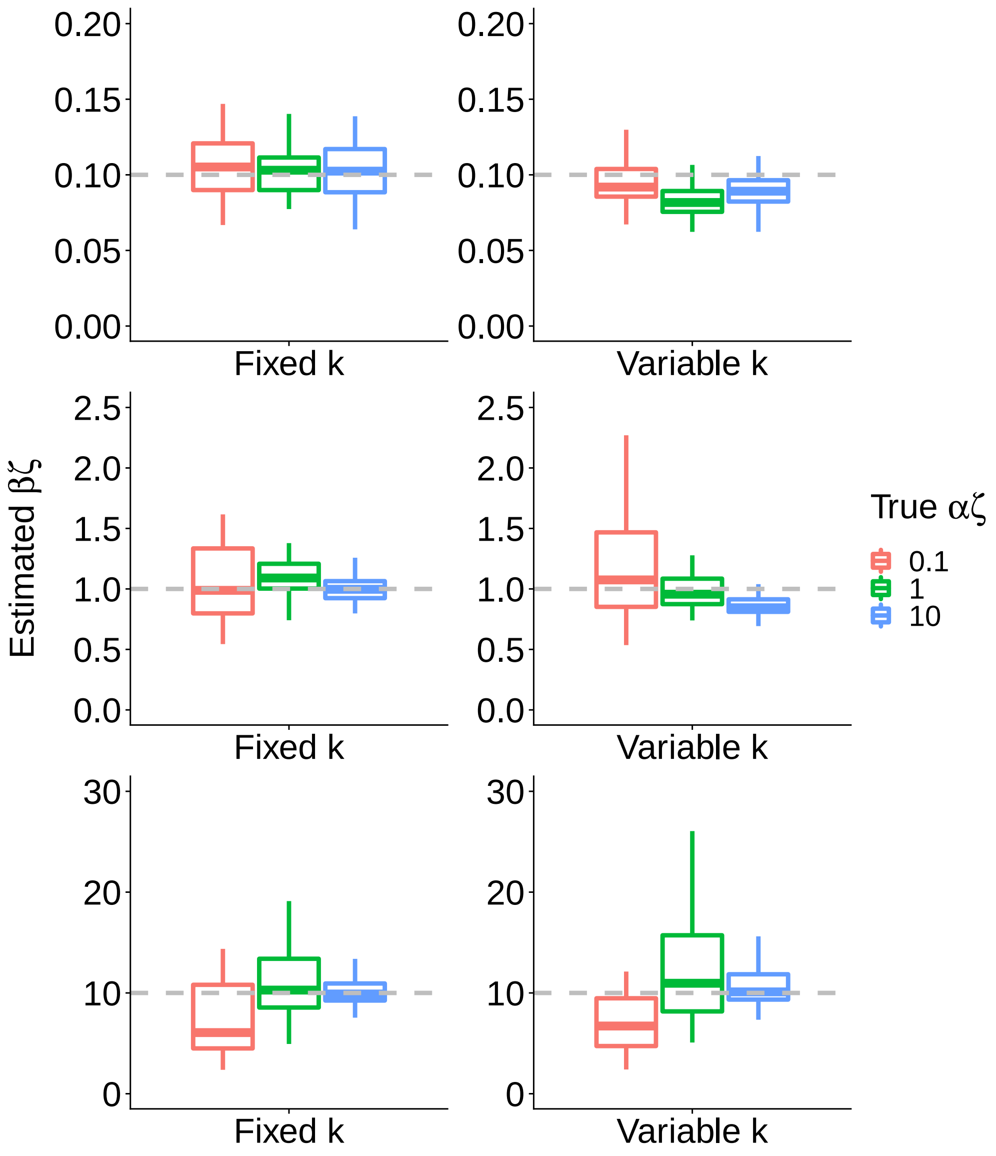


Supplementary Figure S2: Estimated values of $\beta\zeta$ under the variable-pause-site model for simulated true values of $\beta\zeta \in \{0.1, 1.0, 10\}$ (top to bottom) and $\alpha\zeta \in \{0.1, 1, 10\}$ (see key), when the pause-site $k$ is fixed (left) or variable across cells (right) in simulation. This is an expanded version of **Fig. 3A** showing a broader range of $\beta\zeta$ values. As in that case, dashed lines indicate the ground truth; boxplots summarize 50 replicates of the simulation; box boundaries indicate 1st and 3rd quartiles, and horizontal line indicates median. A value of $\zeta = 2$ kb/min is assumed so that $\alpha\zeta$ and $\beta\zeta$ can be assumed to have units of events per minute. Pause sites occur at a mean position of $k = 50$ bp. In the variable case, we assume a Gaussian distribution with a standard deviation of 25 bp.

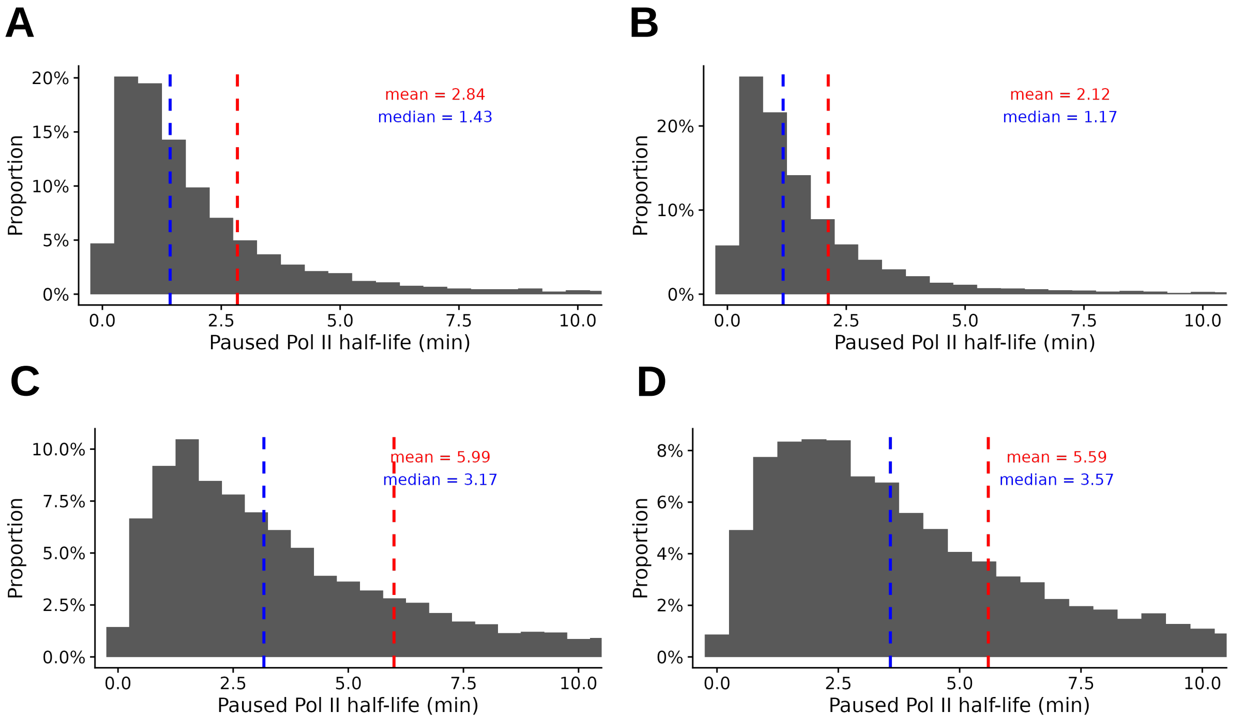


Supplementary Figure S3: Distributions of half-lives for paused RNAPs for (A) the untreated K562 cells from Vihervaara et al. [1], (B) the untreated K562 cells from Dukler et al. [2], (C) the heat-shock-treated K562 cells from Vihervaara et al. [1], and (D) the celastrol-treated (160-min) K562 cells from Dukler et al. [2]. The histograms are truncated at 10 min. for clarity. See **Methods** for details on the calculation of half-lives from $\beta$ estimates.

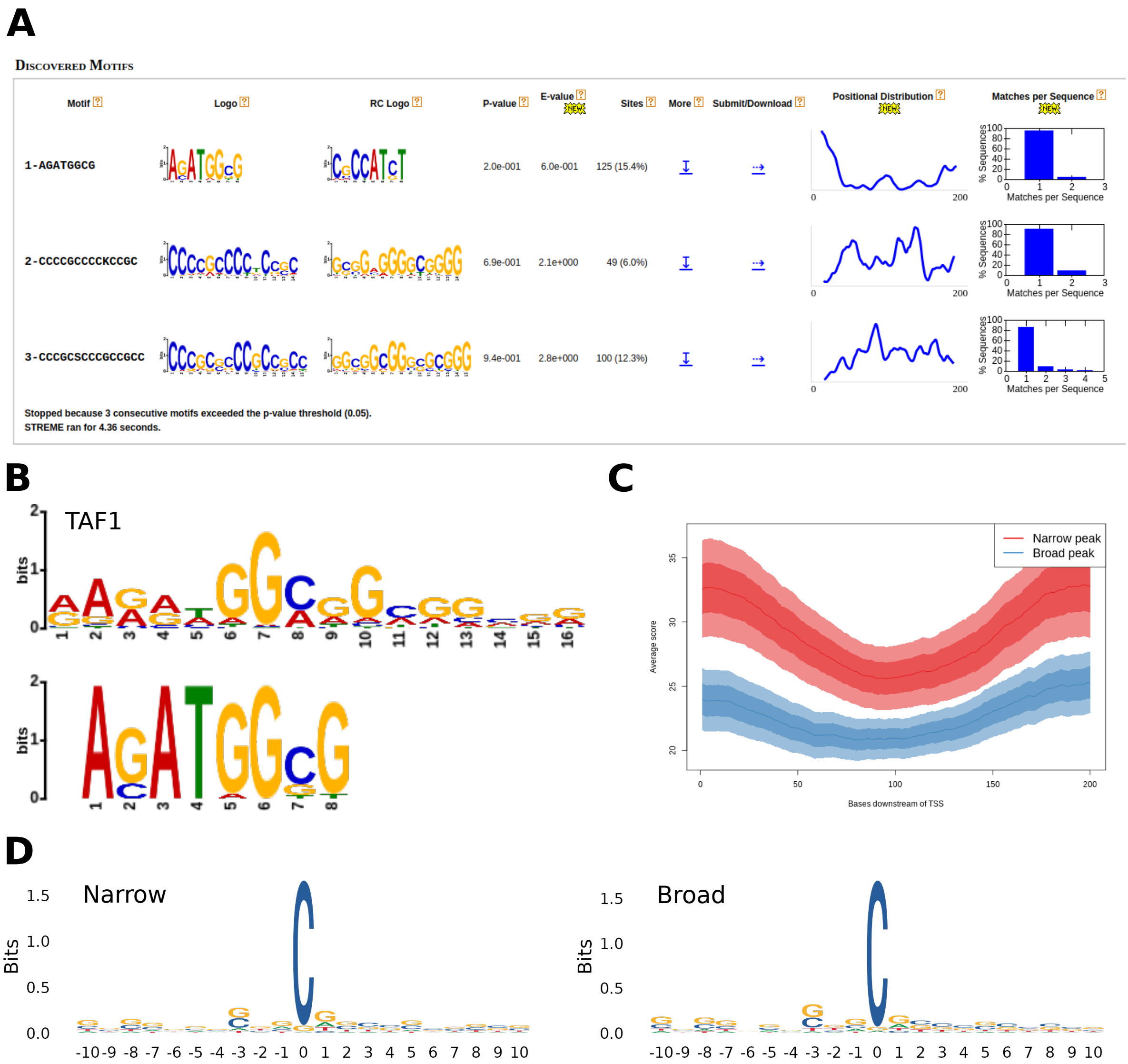


Supplementary Figure S4: Motifs identified in pause peaks from the control sample from the heat-shock data set [1] **A.** Motifs enriched in the 10% genes with narrowest pause peaks (smallest $\hat{\sigma}^2$) compared with the 10% of genes with broadest peaks (largest $\hat{\sigma}^2$) as identified by STREME [3]. **B.** The top candidate from STREME matches the binding motif of TAF1 [4]. **C.** TAF1 ChIP-seq signals from ENCODE [5] for narrow (red) and broad (blue) peaks downstream of the TSS. **D.** Sequences enriched at the site having the maximum PRO-seq read count in the pause peak (200 bp downstream of the TSS).

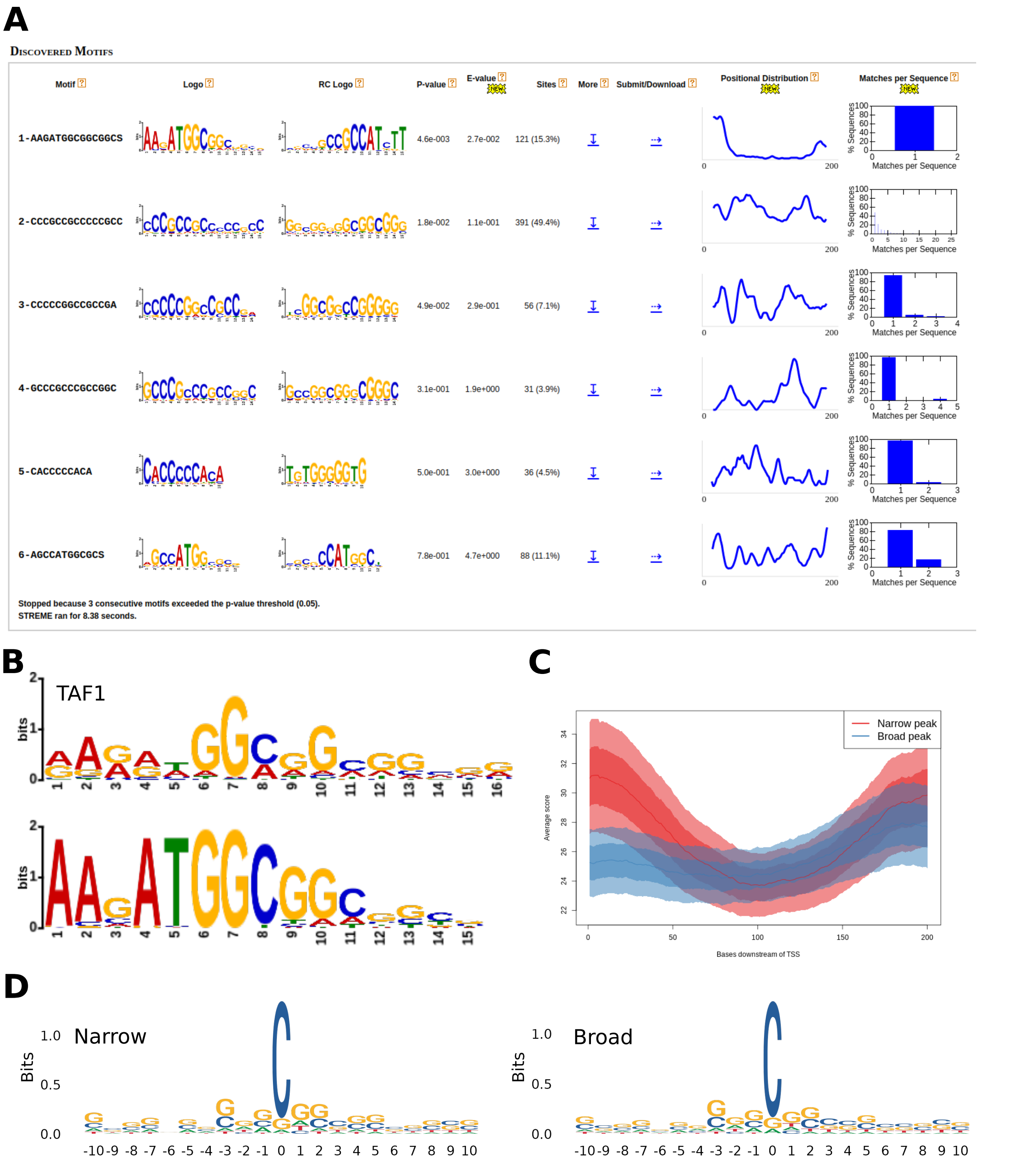


Supplementary Figure S5: Motifs identified in pause peaks from control sample from the celastrol data set [2] **A.** Motifs enriched in the 10% genes with narrowest pause peaks (smallest $\hat{\sigma}^2$) compared with the 10% of genes with broadest peaks (largest $\hat{\sigma}^2$) as identified by STREME [3]. **B.** The top candidate from STREME matches the binding motif of TAF1 [4]. **C.** TAF1 ChIP-seq signals from ENCODE [5] for narrow (red) and broad (blue) peaks downstream of the TSS. **D.** Sequences enriched at the site having the maximum PRO-seq read count in the pause peak (200 bp downstream of the TSS).

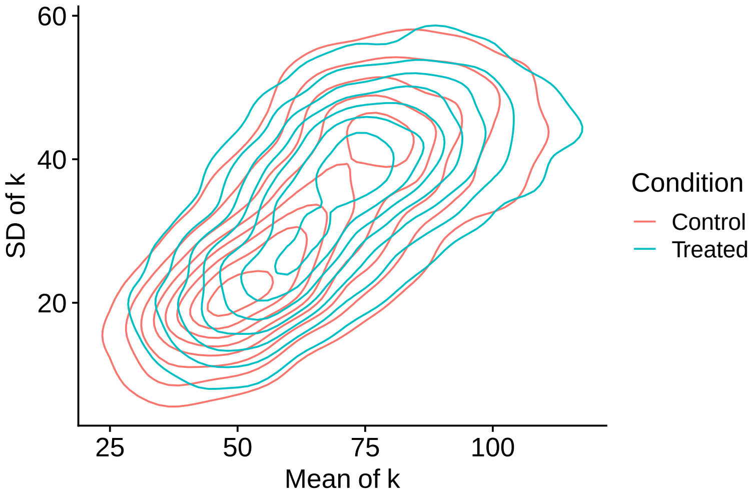


Supplementary Figure S6: Contour plot showing the distribution of estimated means (horizontal axis) and standard deviations (vertical axis) of the pause peak position $k$, under the Celastrol (“Treated”) and Control conditions. Data from ref. [2]. Compare with **Fig. 3C**

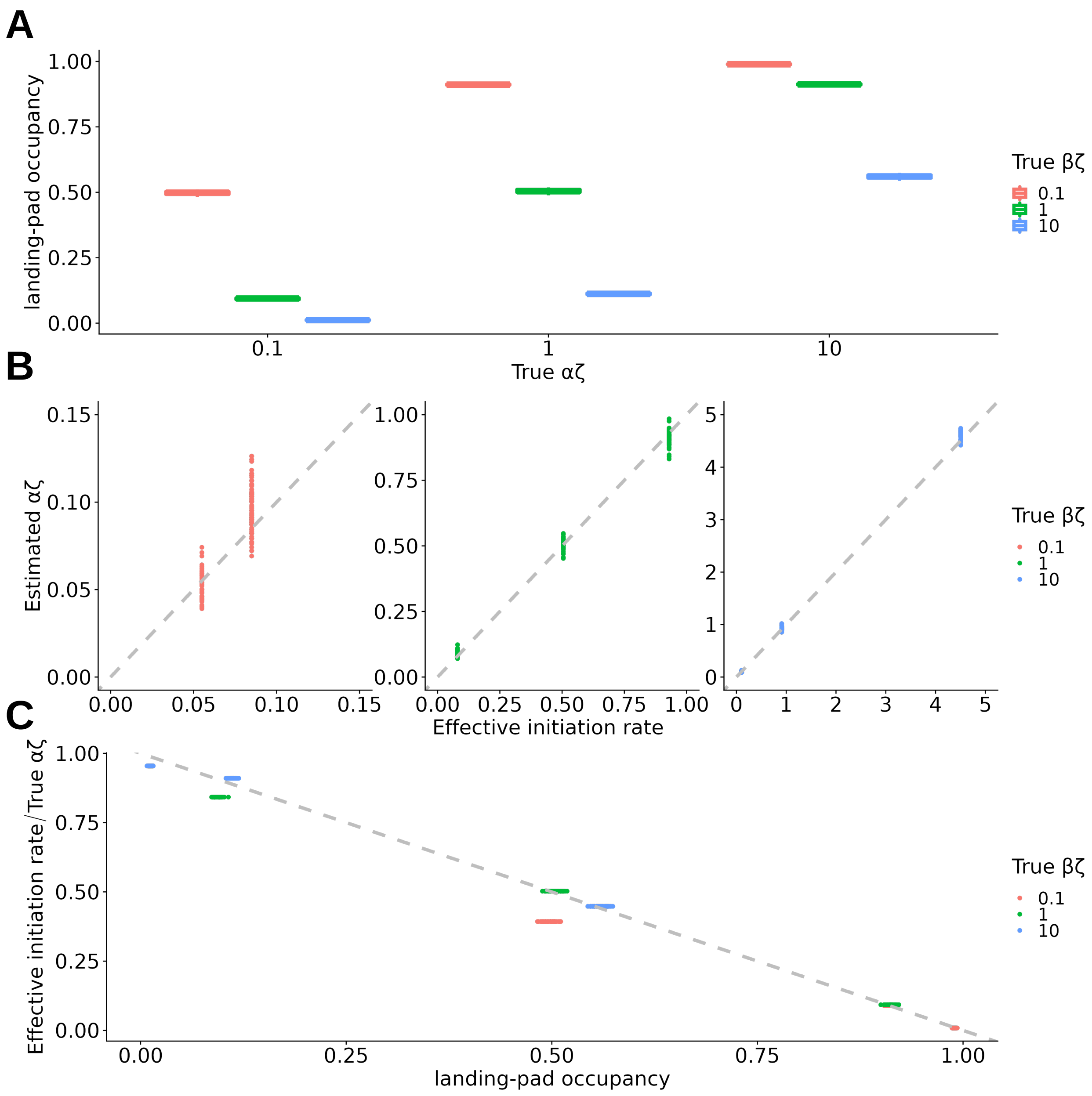


Supplementary Figure S7: Auxiliary data from simulations revealing prevalance of steric hindrance. **A.** Fraction of cells in which the "landing pad" (here, the first 50 bp) for a potential new initiation event is already occupied by an RNAP, for $\alpha\zeta \in \{0.1, 1, 10\}$ (left to right) and $\beta\zeta \in \{0.1, 1, 10\}$ (see key). **B.** Rates at which initiation rates successfully occur ("effective initiation rate") vs. estimates of $\alpha\zeta$, for true $\alpha\zeta \in \{0.1, 1, 10\}$ (left to right) and $\beta\zeta \in \{0.1, 1, 10\}$ (see key). The estimates of $\alpha\zeta$ are much closer to the effective initiation rates than to the true initiation rates. **C.** Landing-pad occupancy (as in panel **A**) vs. "correctness" of estimated $\alpha\zeta$, as measured by the ratio of the estimated value to the true value. The correctness decreases approximately linearly with the landing-pad occupancy.

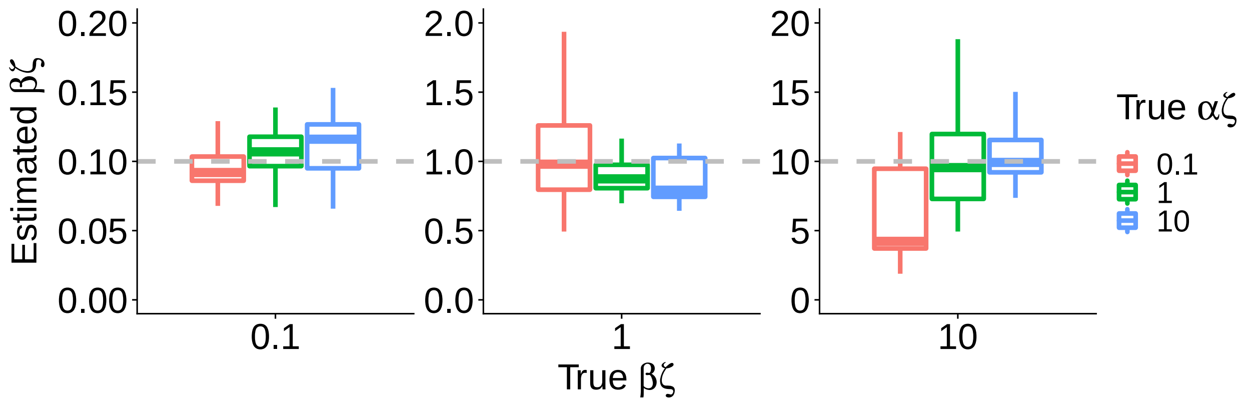


Supplementary Figure S8: Estimated values of $\beta\zeta$ under the steric hindrance model for simulated true values of $\beta\zeta \in \{0.1, 1.0, 10\}$ (left to right) and $\alpha\zeta \in \{0.1, 1, 10\}$ (see key). Estimates are close to the truth, except for the case of $\beta\zeta = 10$, $\alpha\zeta = 0.1$, for which the pause peak was poorly defined. As in previous plots, dashed lines indicate the ground truth; boxplots summarize 50 replicates of the simulation; box boundaries indicate 1st and 3rd quartiles, and horizontal line indicates median. A value of $\zeta = 2$ kb/min is assumed so that $\alpha\zeta$ and $\beta\zeta$ can be assumed to have units of events per minute. Pause sites are variable with a mean position of $k = 50$ bp and a standard deviation of 25 bp.

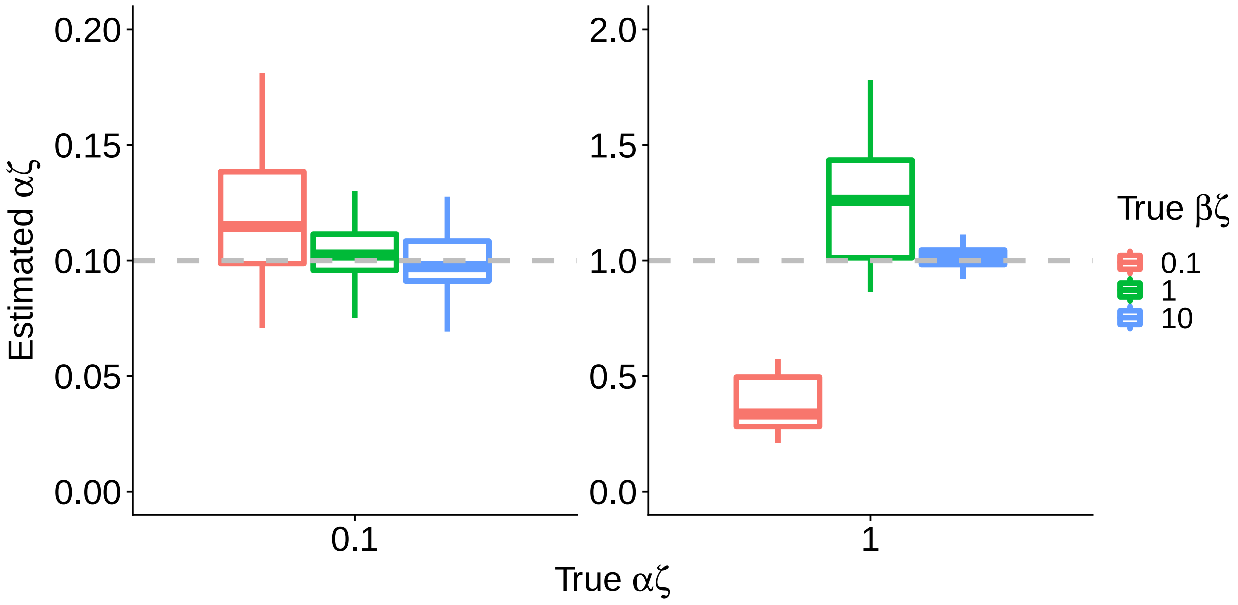


Supplementary Figure S9: Estimated values of $\alpha\zeta$ under the steric hindrance model, for $\alpha\zeta \in \{0.1, 1\}$ (left to right) and $\beta\zeta \in \{0.1, 1, 10\}$ (see key). The poor estimate for $\alpha\zeta = 1, \beta\zeta = 0.1$ reflects underestimation of $\phi$ near the boundary of $\phi = 1$. For similar reasons, estimates with $\alpha\zeta = 10$ were highly variable and therefore are omitted. As in previous plots, dashed lines indicate the ground truth; boxplots summarize 50 replicates of the simulation; box boundaries indicate 1st and 3rd quartiles, and horizontal line indicates median. A value of $\zeta = 2$ kb/min is assumed so that $\alpha\zeta$ and $\beta\zeta$ can be assumed to have units of events per minute. Pause sites are variable with a mean position of $k = 50$ bp and a standard deviation of 25 bp.

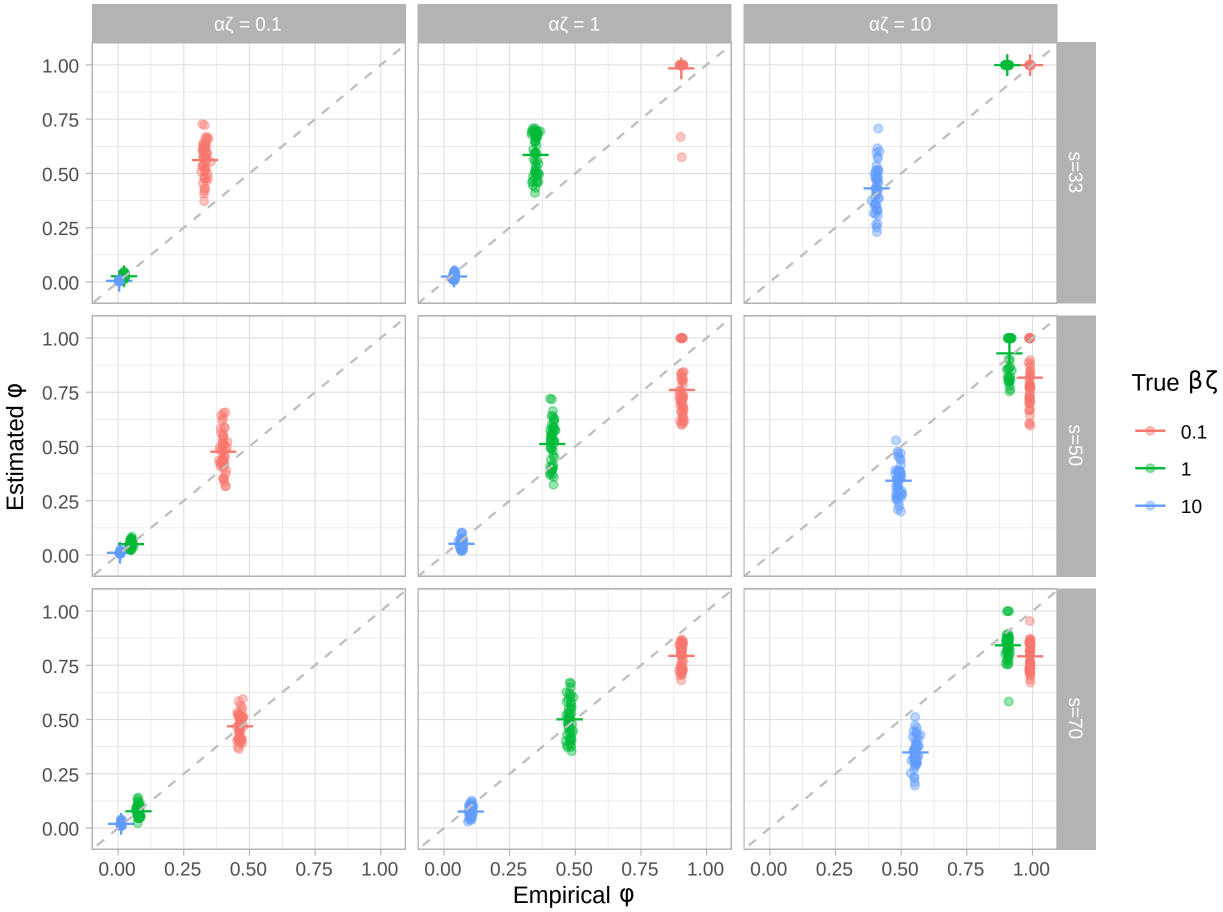


Supplementary Figure S10: Accuracy of estimated landing-pad occupancy $\phi$ under the steric hindrance model, for various values of $\alpha\zeta$ (columns), the spacing $s$ between RNAPs (rows), and $\beta\zeta$ (see key). Scatter plots show the fraction of simulated cells for which the first 50 bp (the "landing-pad") are occupied by an RNAP at steady state ("Empirical $\phi$") vs. the fraction predicted to be occupied under the model ("Estimated $\phi$") based on the simulated NRS data. 50 simulations were performed per parameter combination. Dashed line indicates $y = x$, and colored crosses represent the means of the corresponding points. A value of $\zeta = 2$ kb/min is assumed, so that $\alpha\zeta$ and $\beta\zeta$ are in events per minute.

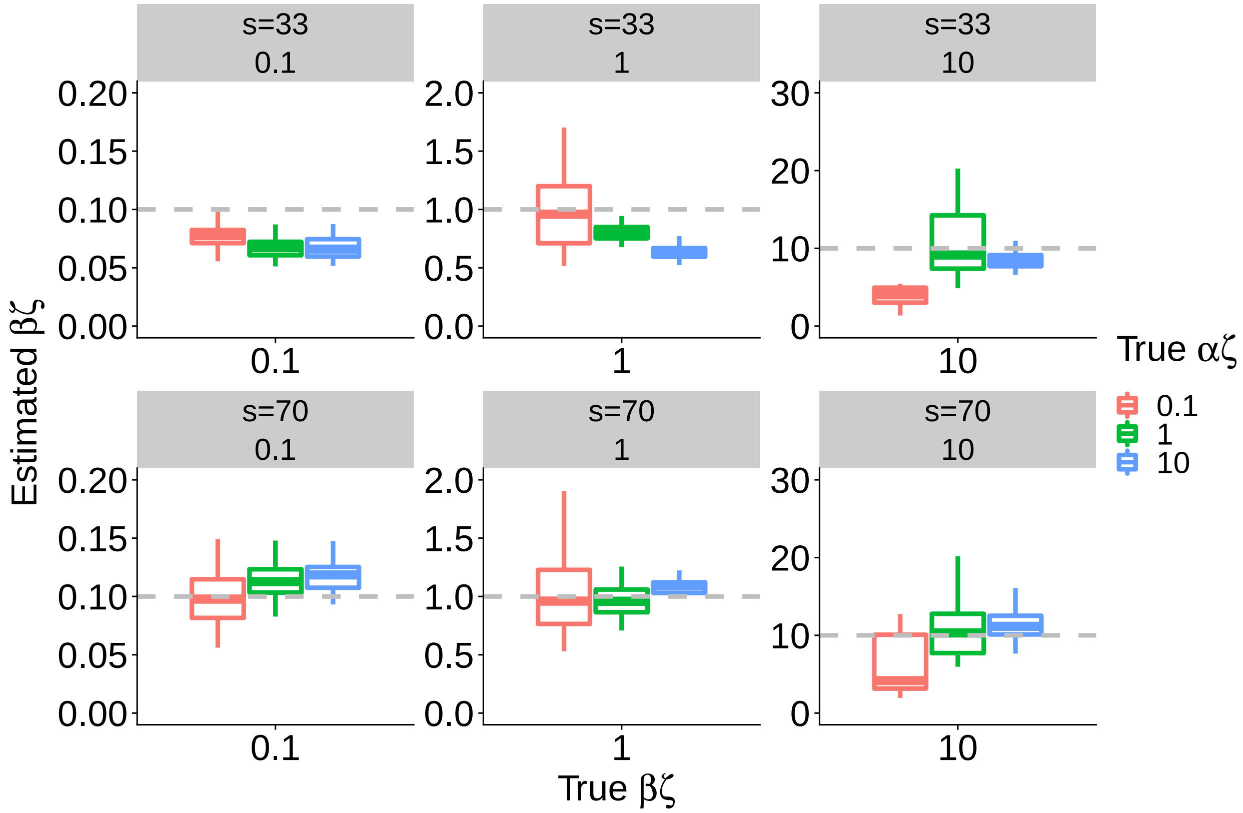


Supplementary Figure S11: Estimated values of $\beta\zeta$ under the steric hindrance model with alternative choices of the spacing parameter $s$ (rows). Results are shown for simulated true values of $\beta\zeta \in \{0.1, 1.0, 10\}$ (left to right) and $\alpha\zeta \in \{0.1, 1, 10\}$ (see key). As in previous plots, dashed lines indicate the ground truth; boxplots summarize 50 replicates of the simulation; box boundaries indicate 1st and 3rd quartiles, and horizontal line indicates median. A value of $\zeta = 2$ kb/min is assumed so that $\alpha\zeta$ and $\beta\zeta$ can be assumed to have units of events per minute. Pause sites are variable with a mean position of $k = 50$ bp and a standard deviation of 25 bp.

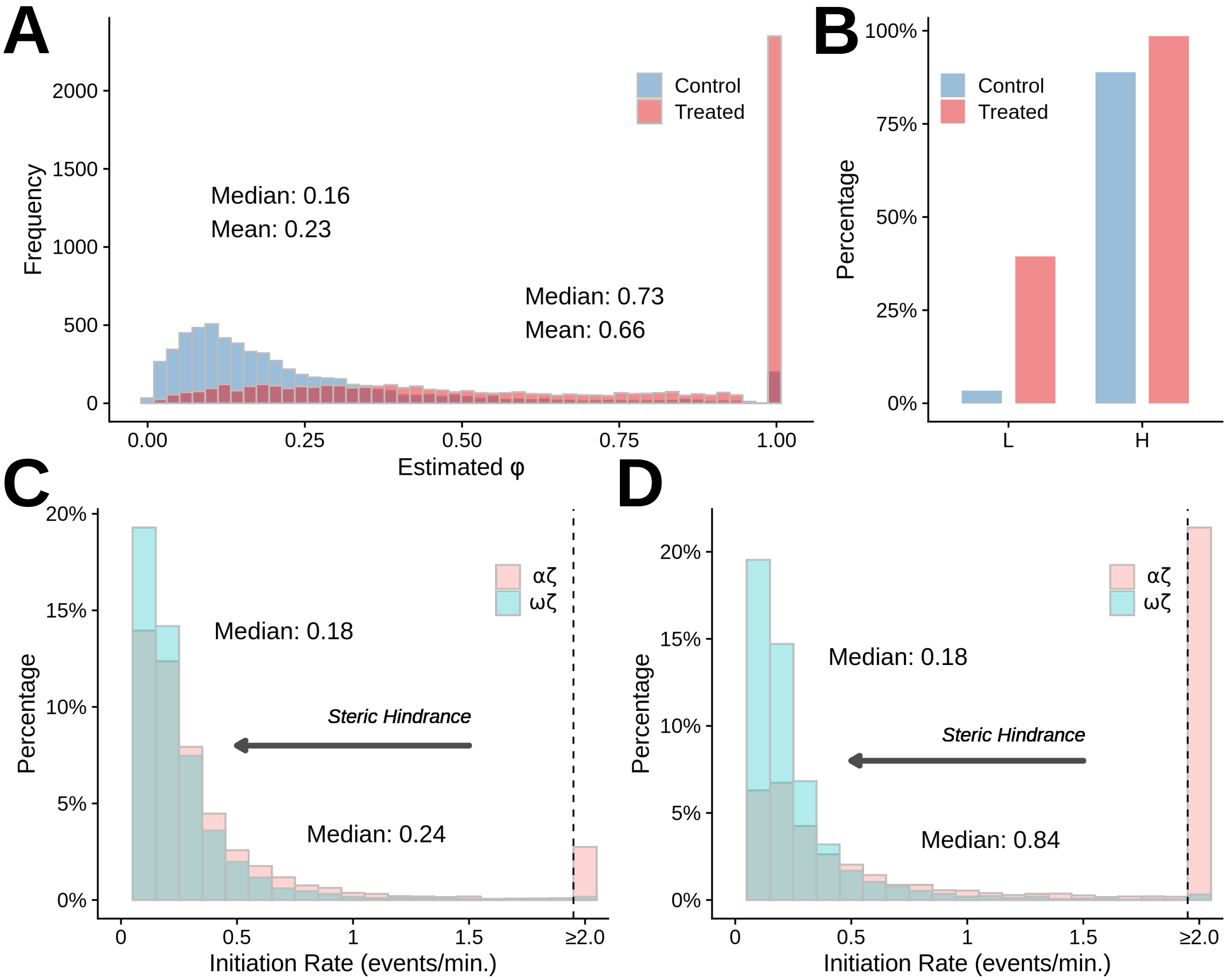


Supplementary Figure S12: **A.** Distribution of estimated $\phi$ for 5,964 robustly expressed genes in K562 cells before (Control) and after (Treated) treatment with celastrol under the low (L) calibration [2] (see **Methods** for details). **B.** Percentages of genes having fully occupied landing-pads ($\phi \approx 1$) before (Control) and after (Treated) treatment with celastrol, under the low (L) and high (H) calibrations. **C & D.** Distributions of scaled estimates of the "effective" ($\omega\zeta$) and "potential" ($\alpha\zeta$) rates of transcription initiation, in events per minute per cell, for the same genes. Panel **C** represents the NHS case and panel **D** represents the HS case. The $x$-axes are truncated to highlight the bulk of the distributions. Gray arrows indicate effects of steric hindrance.
